# Unconventional components complement the cryptic kinetochore of the ciliate *Tetrahymena thermophila*

**DOI:** 10.1101/2025.11.28.691167

**Authors:** Emine I. Ali, Maximilian W. D. Raas, Laura E. van Rooijen, Paula Sobrevals Alcaraz, Harmjan R. Vos, Eelco C. Tromer, Berend Snel, Geert J.P.L. Kops

**Affiliations:** Oncode Institute, Utrecht, the Netherlands; Hubrecht Institute, Royal Netherlands Academy of Arts and Sciences (KNAW), Utrecht, the Netherlands; University Medical Center Utrecht, Utrecht, the Netherlands; Theoretical Biology & Bioinformatics, Department of Biology, Faculty of Science, Utrecht University, Utrecht, the Netherlands; Center for Molecular Medicine, Molecular Cancer Research, University Medical Center Utrecht, Utrecht, the Netherlands; Cell Biochemistry, Groningen Biomolecular Sciences and Biotechnology Institute, Faculty of Science and Engineering, University of Groningen, Groningen, the Netherlands

## Abstract

The accurate segregation of chromosomes is mediated by kinetochores, multi-protein structures that connect centromeric chromatin to the dynamic microtubules of the spindle apparatus. Comparative genomics surveys predict a complex kinetochore in the last eukaryotic common ancestor (LECA) and recurrent loss or replacement of its subcomplexes across the eukaryotic tree of life. Understanding kinetochore composition and organization in diverse lineages can reveal the trajectories of kinetochore evolution in eukaryotes and aid in dissecting the function of each subcomplex. *Tetrahymena thermophila* is a unicellular eukaryote of the phylum Ciliophora with a largely elusive kinetochore composition. Here, we leverage proximity proteomics coupled to deep homology detection approaches to identify 16 kinetochore proteins in *T. thermophila*, dubbed KiTTs (**<u>Ki</u>**netochore of ***<u>T</u>****etrahymena **<u>t</u>**hermophila* 1-16). We find that nine KiTTs (3-9 + 15-16) are cryptic orthologs of conventional kinetochore proteins that previously remained undetected due to extensive sequence divergence. Four KiTTs (10-13) are not orthologous to known subunits and therefore represent unconventional kinetochore proteins. Super-resolution imaging places three of these novel proteins (KiTT10/11/13) at the inner kinetochore, whereas the fourth (KiTT12) localizes near the MIS12 complex at the outer kinetochore. RNAi-mediated depletion of KiTT12 reduces levels of the outer kinetochore protein KiTT1^NDC80^ and causes chromosome segregation errors, showcasing a bona fide role at the kinetochore. Our work reveals a unique kinetochore composition in a ciliate, providing new insights into the evolution of an essential cellular protein machine.

## INTRODUCTION

Kinetochores are multi-protein structures that mediate the accurate attachment of chromosomes to the spindle microtubules during cell division [1–3]. In the majority of eukaryotes, kinetochores bind microtubules via the KMN network, consisting of the KNL1, MIS12, and NDC80 complexes. The NDC80 complex forms the main microtubule binding interface and together with additional protein complexes like SKA (e.g. in animals) or the analogous Dam complex (e.g. in fungi) tracks dynamic microtubules [4–6]. The KNL1 complex can also bind microtubules and scaffolds error correction and spindle assembly checkpoint (SAC) machinery [7]. The MIS12 complex links the NDC80 and KNL1 complexes to chromatin via a network of proteins known as the constitutive centromere-associated network (CCAN). In human cells this network consists of 16 proteins assembled into five subcomplexes [1,8,9]. Of these, CENP-C and CENP-T directly link the KMN network to the centromeric chromatin, epigenetically defined by nucleosomes containing the centromere-specific histone H3 variant CENP-A [8,10–13]. The majority of our knowledge of kinetochore organization and function stems from studies using yeast and animal cells, representing only a small fraction of eukaryotic diversity (**Figure 1A**) [14]. As a result, we have a limited understanding about kinetochore composition and function from other branches of the eukaryotic tree of life.

**Figure 1.**
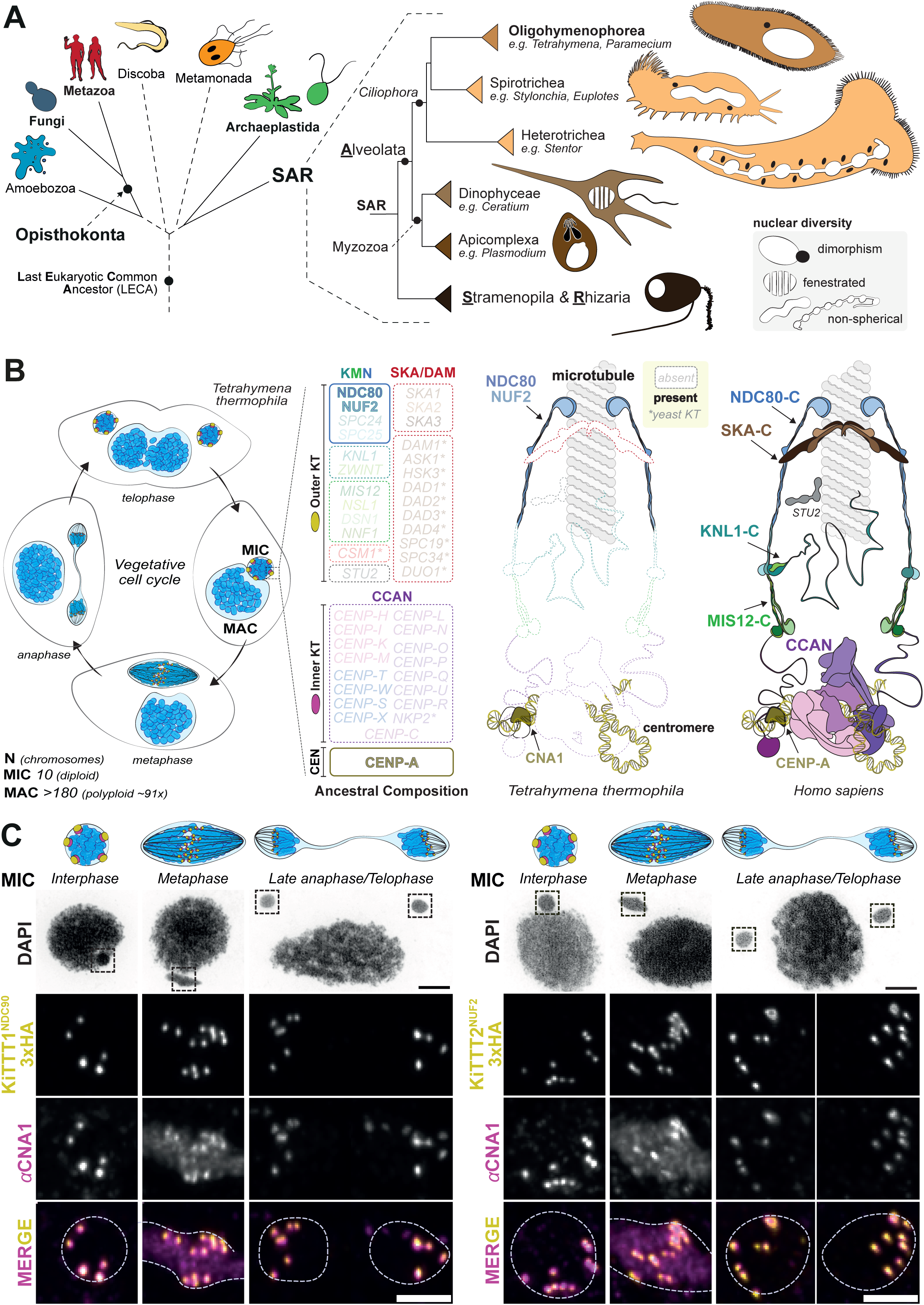
Tetrahymena thermophila kinetochore proteins localise constitutively to centromeres. (A) Phylogenetic position of *Tetrahymena thermophila* in the eukaryotic tree of life. Representative organisms for various (super)groups; silhouettes from left to right: *Dictyostelium discoideum*; *Saccharomyces cerevisiae*; *Homo sapiens*; *Trypanosoma brucei*; *Giardia intestinalis*; *Arabidopsis thaliana*; *Chlamydomonas reinhardtii*. Right: increasing zoom in on the tree for *Tetrahymena* within the supergroup SAR (Stramenopila-Alveolata-Rhizaria), the infrakingdom Alveolata and lastly the class Oligophymenophorea. Remarkably, alveolates showcase extensive morphological and nuclear diversity: (1) nuclear dimorphism and strange nuclear shapes for the macronucleus amongst ciliates; (2) fenestrated nuclei amongst dinoflagellates where microtubules of the spindle apparatus that run through the nucleus during mitosis and connect to chromosomes through the nuclear envelope; and a specialised nucleus area that houses clustered centromeres near the microtubule organising centre (MTOC) [27,31–33]. (B) <u>Left:</u> vegetative life cycle of *T. thermophila.* Each cell contains two distinct nuclei; a germline micronucleus (MIC) and a somatic macronucleus (MAC). The MIC segregates mitotically and MAC chromosomes segregate amitotically. <u>Right:</u> schematic depiction of the predicted kinetochore composition in *T. thermophila* and *H. sapiens* relative to the inferred composition in the ancestral eukaryote (LECA) [16]. The core conventional kinetochore consists of the inner and outer kinetochore units colored in yellow and magentata. The predicted *T. thermophila* kinetochore are NDC80/NUF2 and CNA1 *ancestral subunit not found at the human kinetochore. (C) The subcellular localisation of KiTT1^NDC80^, KiTT2^NUF2^ and CNA1 during the micronuclear mitosis. Cells expressing KiTT1^NDC80^-3xHA-TurboID (left) and KiTT2^NUF2^-3xHA-TurboID (right) were fixed and stained with antibodies against HA and CNA1. Insets show magnified micronuclei. Scale bars DAPI image: 5 µm, MIC zoom-ins: 2 µm.

Recent comparative surveys of diverse eukaryotic genomes and experimental work in a set of divergent lineages revealed that the complex kinetochore known from yeast and animal studies was already present in the last eukaryotic common ancestor (LECA). However, these surveys also uncovered a surprisingly extensive diversity of kinetochore composition across eukaryotes, despite its ancient origins and essential role in facilitating cell division [15,16]. While CENP-C, CENP-A, and the KMN network, are generally retained, the CCAN is often absent [15], as exemplified in, for example, *Caenorhabditis elegans*, *Drosophila melanogaster* [17,18] and the model plant *Arabidopsis thaliana* [19]. In the basidiomycete yeast *Cryptococcus neoformans*, the CCAN is similarly absent, and its functions seems to have been partly assumed by the protein bridgin [20]. More dramatically, some unicellular flagellates show an apparent complete absence of the conventional kinetochore [21,22], strongly suggesting the presence of evolutionary novel and unconventional kinetochore systems. Indeed, in one such lineage, the kinetoplastids, the conventional kinetochore has been supplanted by a network of >30 proteins [23,24]. Although some components show putative homology to conventional subunits [24,25], most are unique to kinetoplastids.

Cryptic kinetochores can also be found in Apicomplexa, a mostly parasitic clade of which various lineages have highly unconventional nuclear biology and cell division strategies [26,27]. Our previous genome surveys suggested that apicomplexans lacked nearly all conventional kinetochore subunits, with only a handful of candidates such as CENP-A and members of the NDC80 complex detectable [15]. However, subsequent integrative work combining targeted proteomics with highly sensitive bioinformatic searches in the “grey zone” of detection —including profile Hidden Markov Models (HMMs) and 3D structure comparisons with AlphaFold— revealed that many bona fide conventional kinetochore proteins are in fact present, having extremely divergent amino acid sequences [28,29]. In the malaria parasite *Plasmodium* for instance, a cryptic KMN network was found to attach to multiple CENP-C orthologs at the inner kinetochore supplemented by lineage-specific subunits, illustrating how previously thought canonical kinetochore protein absences instead reflect amino sequence divergences amongst apicomplexans [28]. The discovery of cryptic and analogous kinetochores throughout the eukaryotic diversity highlights both the evolutionary plasticity of these structures and the technical challenges of identifying their components, which often require a combination of experimental validation and advanced homology detection [22,28,30]. Together, these findings underscore the need to explore kinetochore composition and function in evolutionarily distant lineages, particularly those in which only a handful of components have been identified.

*Tetrahymena thermophila* is a unicellular eukaryote that belongs to the phylum Ciliophora (commonly referred to as ciliates) which together with Apicomplexa and Dinoflagellata comprises the Alveolata, a superphylum harboring remarkable variation in their nuclear biology impacting their chromosome segregation strategies [27,31–33] (**Figure 1A**). Ciliates including *T. thermophila* exhibit nuclear dimorphism, with a micronucleus (MIC), functioning as the *de facto* germline, and a somatic macronucleus (MAC) which is the location of active transcription (**Figure 1A-B**). The MIC contains five diploid chromosomes while the MAC harbors a highly fragmented polyploid version of the genome, with ∼91 copies of 181 transcriptionally active chromosomes which lack centromeres [34]. In mitosis, the MIC undergoes closed mitosis, while the MAC separates into two via amitotic division. The centromeres of MIC chromosomes are defined by the CENP-A ortholog named CNA1 and kinetochores harbor the outer kinetochore proteins NDC80 and NUF2 [35–37]. The genomes of *T. thermophila* and nine other *Tetrahymena* species have been sequenced, allowing for deep genomic sampling of this lineage, which aids in remote homology detection [37–39]. A wide range of genetic manipulation techniques have been used to explore various cellular features of *T. thermophila* in fields of cilia, cell patterning, genome rearrangement, and meiosis [40–42].

The apparent absence of the vast majority of conventional kinetochore components amongst ciliates, and specifically *T. thermophila*, raises the question how CNA1 at the centromere is connected to the outer kinetochore components NDC80 and NUF2, and how this kinetochore evolved. In this study, we combine the sensitive proximity proteomics tool TurboID [43] and lineage-specific deep homology detection protocols to reveal that the *T. thermophila* kinetochore is a hybrid structure made up of cryptic conventional kinetochore components that previously evaded bioinformatic detection, supplemented with four newly-described unconventional kinetochore components of diverse evolutionary origins.

## RESULTS

### Tetrahymena thermophila NDC80 and NUF2 are bona fide kinetochore proteins

To elucidate the kinetochore composition of *T. thermophila*, we first verified the kinetochore localisation of the previously-identified orthologs of NDC80 and NUF2 [15]. We expressed 3xHA-TurboID tagged *Tt*NUF2 and *Tt*NDC80 and performed immunolocalization together with the CENP-A ortholog CNA1. Consistent with the *Tt*NDC80 localisation pattern during meiosis [44], we found that both the orthologs of *Tt*NUF2 and *Tt*NDC80 are restricted to the micronuclei and co-localised with CNA1 signal during mitosis. Furthermore, these proteins localise at kinetochores across the cell cycle (**Figure 1C**). Therefore, localisation of *Tt*NUF2 and *Tt*NDC80 is consistent with a function as *bona fide* kinetochore components and from here-on, we will refer to them as **<u>Ki</u>**netochore of **<u>T</u>**etrahymena **<u>T</u>**hermophila proteins 1 and 2 (KiTT1^NDC80^ and KiTT2^NUF2^).

### Comprehensive identification of Tetrahymena thermophila kinetochore components

Typically, NDC80 and NUF2 dimerise and form a complex together with a dimer of SPC24 and SPC25, together forming the Ndc80 complex [45]. However, orthologs of SPC24 and SPC25 were not found in previous kinetochore comparative genomics efforts for *Tetrahymena* [15]. The availability of genomic sequences of multiple *Tetrahymena* species supplemented with members from Alveolates facilitated highly-sensitive homology detection strategies, including iterative lineage-specific Hidden Markov Models (HMM) searches and HMM-vs-HMM profile searches using deep-sampled lineage-specific profiles [22,29,46]. We performed such sensitive HMM searches against the predicted *T. thermophila* proteome and identified two genes (TTHERM_00312495 and TTHERM_01142690) with putative homology to SPC24 (E=13) and SPC25 (E=0.045), respectively (**Table S1**). Co-folded structures of putative dimer partners within the *T. thermophila* NDC80 contained domains with strong similarity to the known structure found in *Saccharomyces cerevisiae* (**Figure 2A**), supporting their proposed orthology and suggesting they are indeed part of a functional NDC80 complex. To validate the localisation of these proteins to the *T. thermophila* kinetochore, we created endogenous fusion proteins with 3xHA tags and performed immunostainings against HA in combination with a custom antibody against KiTT1^NDC80^ (**Figure S1A**). Both TTHERM_00312495 and TTHERM_01142690 consistently co-localised with KiTT1^NDC80^ in the MIC (**Figure S2**). Thus, we conclude that TTHERM_000312495 and TTHERM_01142690 are indeed *bona fide T. thermophila* orthologs of SPC24 and SPC25, completing the NDC80 complex in the *T. thermophila* kinetochore. From here-on these proteins will be referred to as KiTT3^SPC24^ and KiTT4^SPC25^.

**Figure 2.**
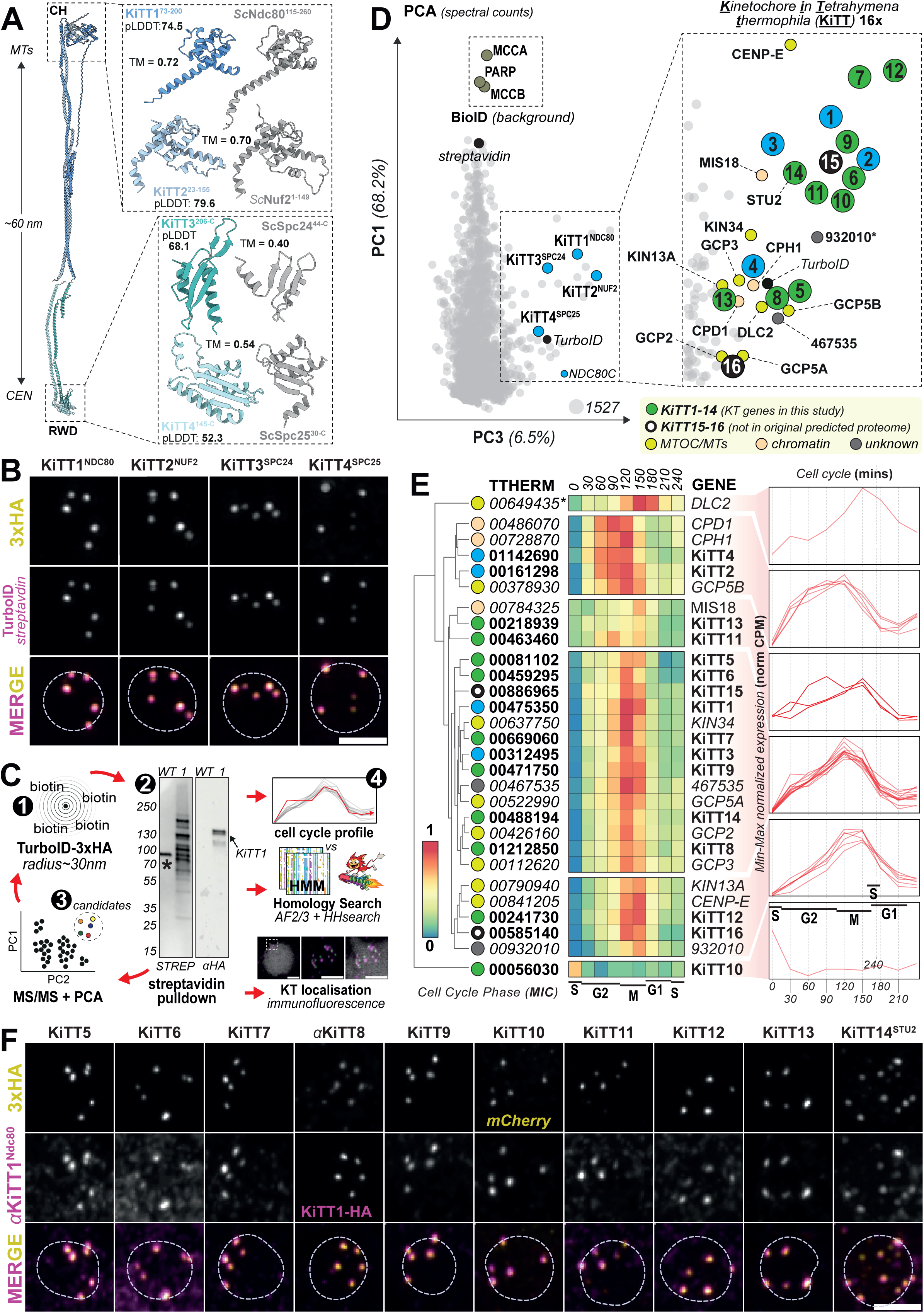
Identification of kinetochore proteins in T. thermophila. (A) AlphaFold 2/3 predicted multimeric structures of NDC80 complex dimers consisting of KiTT1^NDC80^-KiTT2^NUF2^ and KiTT3^SPC24^-KiTT4^SPC25^. Zoom-ins of the CH domains of the KiTT1^NDC80^-KiTT2^NUF2^ dimer are shown side-by-side with the CH domains of the structure of the *Saccharomyces cerevisiae* NDC80/NUF2 dimer (PDB ID: 5TCS)[122]. (B) Immunostaining of interphase MICs with a conjugated streptavidin against biotin (yellow) an antibody against HA for KiTT1^NDC80^/KiTT2^NUF2^/KiTT3^SPC24^/KiTT4^SPC25^-3xHA-TurboID (magenta). Scale bar, 2 µm. (C) Workflow for candidate kinetochore protein detection in four steps (1) BioID is performed using the efficient promiscuous biotinylase TurboID [43]; (2) biotinylated proteins are pulldown using magnetic streptavidin beads - example shows output for BioID for KiTT1^NDC80^ (**Figure S3**); (3) trypsin-cleaved peptides are submitted for LC-MS/MS, and candidates are identified based on Principal components analysis (PCA) of multiple BioID experiments with wildtype cells as controls; 4) candidate selection is further scrutinized based on cell cycle expression profiles, deep homology detection pipelines and kinetochore co-localisation in immunofluorescence assay. This pipeline works in an iterative manner were in step three novel candidates are tagged with TurboID and subsequently used for additional BioID experiments. (D) PCA of integrated spectral counts detected in BioID experiments from KiTT1^NDC80^, KiTT2^NUF2^, KiTT4^SPC25^ and wildtype cells as a control. PC1 and PC3 are plotted as they showed a clear kinetochore-based cluster. NDC80 complex proteins (3 are baits in the BioID) are highlighted in blue, characterized KiTTs (5-14) in green and in black KiTT15-16 that were not part of the original proteome that was used to analyse the mass spectra. Other categories have unique colors (MTOC/SPB & chromatin) and unknown proteins are grey. (E) <u>Left:</u> heatmap showing RNA seq expression levels (min-max normalized) of candidates obtained from the PCA analysis in D across the *T. thermophila* cell cycle. Expression level patterns were manually delineated into clusters based on an average linkage tree. <u>Right:</u> shows overlays of line graphs of expression levels across the cell cycle for each cluster. Expression level data was adapted from Bertagna *et al* [51]. (F) Localisation of KiTT1^NDC80^ and KiTT5-KiTT14 in the interphase MICs. Immunofluorescence staining of KiTT1^NDC80^ (magenta), 3xHA-TurboID tagged candidates (yellow), KiTT10-mCherry (yellow). KiTT1^NDC80^-3xHA-TurboID cells were stained with custom antibodies against KiTT8 (yellow) and HA (magenta). Scale bar, 2 µm. POI= protein of interest.

The tags for KiTT1-4 also included the promiscuous biotinylation enzyme TurboID [43], enabling biotin-based proximity proteomics (BioID) to identify additional components of the *T. thermophila* kinetochore. First, we verified TurboID activity by immunostaining with fluorescently labelled streptavidin, which confirmed efficient and specific biotinylation of proteins in vicinity of KiTT1^NDC80^, KiTT2^NUF2^, KiTT3^SPC24^, and KiTT4^SPC25^ (**Figure 2B, Figure S3**). Because centromeric sequences are specifically eliminated from MAC chromosomes and thus lack kinetochores, we modified previously published protocols to enrich specifically for MICs [47]. Biotinylated proteins were then purified from G2-stage MICs using streptavidin and identified by mass spectrometry (**Figure 2C, Figure S3, Table S2**). Principal component analysis (PCA) of the total spectral counts per identified proteins found amongst three separate BioID experiments (wild type versus KiTT1^NDC80^, KiTT2^NUF2^, KiTT4^SPC25^) revealed similar behaviour of 23 proteins to *Tt*NDC80-C members in PC1 and PC3 (accounting for 68.2% and 6.5% of the variance, respectively; **Figure 2D**). 11 out of the 23 putative kinetochore subunits were annotated with *‘unknown function’*, while the remaining 12 could be assigned to two functional categories. The first category contained chromatin-related proteins: subunits of condensin-I (CAPH1, CAPD1) and MIS18, which indeed have been found to localise near centromeric chromatin in *T. thermophila* [44], and other eukaryotes [48,49], respectively. The second category comprised microtubule-organizing centre (MTOC)-related proteins: subunits of the y-TURC complex (GCP2,GCP3,GCP5A/5B), various kinesins (KIN34, CENP-E, KIN13A), the microtubule polymerase STU2, and a dynein light chain (DLC2). The association with MTOC-related proteins is consistent with y-tubulin localization in a dotted pattern on the spindle indicating that the y-TURC complex might reside at kinetochores or at least in close proximity to the MTOC [50]. Finally, cell-cycle-resolved expression analysis [51,52] revealed highly correlated expression patterns—peaking around the G2/M-phase—for KITT1-4 and our 23 candidate kinetochore proteins, supporting their interconnected roles during mitosis, with the exception of TTHERM_0056030 and TTHERM_00649435 (DLC2; **Figure 2E**).

To examine the localisation of the putative kinetochore subunits annotated with ‘unknown function’ as well as the previously-predicted orthologs of CENP-E (TTHERM_000841202)[15] and STU2 (TTHERM_00488194), we tagged them with either 3xHA-TurboID or mCherry at the C-terminus. Tagging was not attempted for TTHERM_0046753 and was unsuccessful for TTHERM_01212850 due to destabilisation of the protein upon tagging. TTHERM_01212850 was thereafter localized using a custom-raised polyclonal antibody (**Figure S1B**). CENP-E-tagged cells showed no signal, for unknown reasons. Out of the 11 proteins for which we could obtain a clear immunostaining signal, 10 consistently co-localised with KiTT1^NDC80^ at the *T. thermophila* kinetochore in both interphase and metaphase (**Figure 2F, Figure S4**). By contrast, TTHERM_00932010 had a more temporally-constricted co-localisation with KiTT1^NDC80^, specifically during telophase (**Figure S5A)**, and we thus did not consider it to be part of the structural kinetochore in *T. thermophila*. Based on their consistent co-localisation with KiTT1^NDC80^ throughout the cell cycle, we consider the remaining 10 proteins to be part of the *T. thermophila* kinetochore and designated them as KiTT5-14, with STU2 renamed to KiTT14^STU2^. Reciprocal BioID experiments with 3xHA-TurboID lines for KiTT5-6-7 and KiTT9 did not reveal any additional kinetochore candidates (**Figure S3, Figure S6, Table S2**).

### KiTT5-9 and KiTT15-16 are cryptic orthologs of conventional kinetochore components

Having identified nine *Tetrahymena* kinetochore proteins with unknown functions in addition to KiTT1-4^NDC80-C^ and KiTT14^STU2^, we next set out to uncover their evolutionary origins. We first investigated whether the KiTTs are highly-divergent orthologs of conventional kinetochore components that previously evaded detection. To do so, we collected the predicted proteomes of 10 *Tetrahymena* species and generated orthologous groups (OGs), allowing for HMM-vs-HMM-based homology detection using the profile HMMs of the KiTTs [38]. Here, we searched against a curated database of full-length and domain-specific profile HMMs of kinetochore proteins [16,22,28] supplemented with OGs from the COG/KOG database [53] (**Figure 3A, Table S1**). To further scrutinize the HMM-vs-HMM hits we performed manual sequence analysis and (multimer) protein structure predictions to corroborate putative homologies. Surprisingly, our approach yielded seven highly divergent homology links for KiTTs with conventional kinetochore proteins (see below). For four KiTTs (KiTT10-13) we could not detect any orthology with canonical kinetochore proteins using our approach and from here-on we refer to them as unconventional kinetochore proteins.

**Figure 3.**
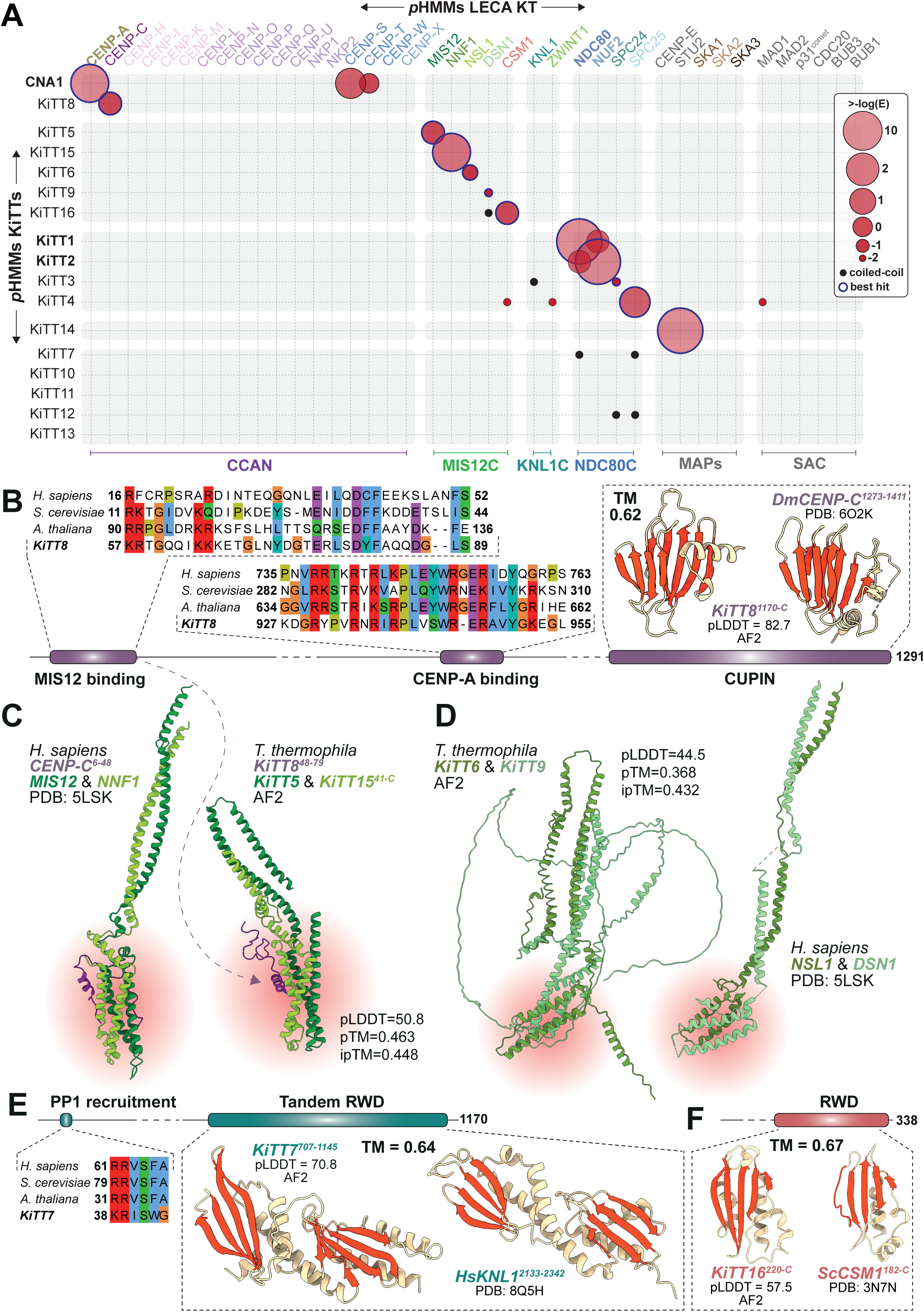
Conventional kinetochore proteins KiTT5-9 & KiTT14-15. (A) HMM-vs-HMM homology search of KiTT profiles versus a curated kinetochore profile database. For abbreviations of complex names see Figure 1B; MAPs: microtubule-associated proteins. (B) Analysis of the KiTT8^CENP-C^ sequence, including putative interaction motifs aligned to known CENP-C orthologs from *Homo sapiens* (human), *Saccharomyces cerevisiae* (budding yeast) and *Arabidopsis thaliana* (thale cress). The C-terminal region was predicted by AF2 to fold into a recognisable CUPIN domain, shown side-by-side to the known structure of the CUPIN domain of the CENP-C ortholog in *Drosophila melanogaster* (fruit fly. (C) Left: human MIS12/NNF1 dimer (greens) with the MIS12-binding motif of CENP-C (purple), PDB ID: 5LSK. Right: Predicted structure of the KiTT5^MIS12^/KiTT15^NNF1^ dimer (greens) together with the putative N-terminal Mis12-binding motif of KiTT8^CENP-C^ (purple). The head domain of the dimer is highlighted in red. (D) Left: Predicted KiTT6^NSL1^/KiTT9^DSN1^ dimer. Right: Human NSL1/DSN1 dimer. PDB ID: 5LSK. The head domain of the dimer is highlighted in red. (**E**) Analysis of the KiTT7^KNL1^ sequence, highlighting the putative PP1 recruitment motif in the N-terminal region. A prediction of the structure of the C-terminal region of KiTT7^KNL1^ with AF2 revealed a tandem RWD domain with strong similarity to the C-terminal tandem RWD of human KNL1 (TM = 0.64, PDB: 8Q5H). (**F**) Analysis of the KiTT16^CSM1^ sequence. A prediction of the structure of the C-terminal region with ESMfold revealed an RWD domain with strong similarity to the C-terminal tandem RWD of *S. cerevisiae* CSM1 (TM = 0.67, PDB: 3N7N).

#### CENP-C

our HMM-vs-HMM approach led to a homology link of KiTT8 (TTHERM_01212850) with the CENP-C CUPIN domain (**Figure 3A**). Structure predictions of the C-terminal region of KiTT8 revealed a fold with significant similarity to the C-terminal CUPIN domain of the CENP-C ortholog characterized in *Drosophila melanogaster* [54](TM score = 0.62, **Figure 3B**). We also identified a motif in KiTT8 with similarity to the CENP-C motif, which typically binds CENP-A [55], as well as a putative MIS12-binding motif in the N-terminal region of KiTT8 [56–59] (**Figure 3B**). Combined, these observations show that KiTT8 is the *T. thermophila* CENP-C ortholog (KiTT8^CENP-C^).

#### MIS12C

our deep homology detection approaches found that the KiTT5 (TTHERM_000081109), KiTT6 (TTHERM_000459299) and KiTT9 (TTHERM_00471750) profile HMMs yielded significant similarity to those of MIS12, NSL1 and DSN1, respectively (**Figure 3A, Table S1**). The fourth MIS12 complex subunit NNF1 was conspicuously absent. Through identification of likely NNF1 orthologs in other *Tetrahymena* species, we tracked down an unannotated genomic region in the *T. thermophila* genome that showed strong sequence similarity to these annotated putative NNF1 orthologs, implying an erroneously-missed prediction of a gene in this region. Indeed, a recent re-annotation of the *T. thermophila* genome includes a gene model matching our predicted NNF1 ortholog (TTHERM_00886965) [60]. Re-mapping peptides from our proximity proteomics data to our predicted NNF1 sequence identified NNF1 in the kinetochore-based TurboID-tagged proteins but not in the wildtype cells (‘15’ in **Figure 2D, Figure S3, Figure S6, Table S2**). Pairwise AF2 co-folding of the predicted structures of the MIS12/NNF1 and NSL1/DSN1 dimers showed a striking similarity to the known structure of the human MIS12 complex, including the characteristic head domains (**Figure 3C-D**). For the MIS12/NNF1 dimer, we also included the putative MIS12C-binding motif of KiTT8^CENP-C^, which was indeed predicted to interact with the MIS12/NNF1 dimer in a similar manner to the human structure (**Figure 3C**) [56]. Taken together, we find strong evidence for the presence of a complete MIS12 complex, composed of KiTT5^MIS12^, KiTT6^NSL1^, KiTT9^DSN1^, and a previously unannotated NNF1 homolog that we hereby designate KiTT15^NNF1^.

#### KNL1 and SAC proteins

KNL1 is characterized by a large disordered region with an N-terminal RVSF motif, responsible for PP1 recruitment [61,62], and a C-terminal tandem RWD domain [63]. Upon close inspection of its sequence, one of our hits obtained from the BioID experiments, KiTT7 (TTHERM_00669060), appeared to harbour an RVSF-like motif (RISW, **Figure 3E**). An AF2 model of the KiTT7 structure revealed the presence of a highly-divergent tandem RWD domain with strong structural similarity to the human KNL1 C-terminal region (TM score = 0.64, **Figure 3E**) [4]. Based on these observations, we consider KiTT7 to be a *T. thermophila* KNL1 ortholog. Interestingly, we have been unable to find any candidate orthologs of KNL1’s dimerization partner ZWINT1, indicating that KiTT7^KNL1^ may have lost this interaction. In addition, we found no evidence of MELT-like motifs that recruit spindle assembly checkpoint (SAC) proteins [64,65] in the KiTT7^KNL1^ sequence. Indeed, amongst the KiTTs we did not find any remote homology with SAC proteins (**Figure 3A, Table S1**). Although not present in any of the BioID experiments, HMM-based searches did identify a MADBUB homolog (TTHERM_00393260) containing only its N-terminal TPR domain, but immunolocalization of the mNeonGreen-tagged protein did not show fluorescent signal (**Figure S5B**).

#### CSM1

inspired by our identification of KiTT15^NNF1^, we explored the possibility of further missing conventional kinetochore proteins not yet annotated in the *T. thermophila* proteome. To do so, we queried all *Tetrahymena* HMM profiles that did not contain a *T. thermophila* ortholog to our curated kinetochore database. This revealed the presence of a CSM1 ortholog (component of the Monopolin complex) conserved across *Tetrahymena* species, but with the curious absence of a *T. thermophila* ortholog. We then again queried the putative *Tetrahymena* CSM1 orthologs against the *T. thermophila* genome, leading to the identification of a previously-unannotated gene (new version proteome: TTHERM_00585140 [60]). Indeed, the obtained gene model showed similarity to the CSM1 RWD HMM model (bitscore = 36.4, **Figure 3A, Table S1**), which was corroborated by an AF2 prediction of a C-terminal RWD domain with similarity to the RWD domain of *Saccharomyces cerevisiae* CSM1 (TM score = 0.67, **Figure 3F**). Re-mapping the sequence of TTHERM_00585140 to peptides from our proximity proteomics showed it to be present in kinetochore-specific BioID experiments (‘16’ in **Figure 2D**, **Figure S3-S6, Table S2**). Hence, TTHERM_00585140 was renamed KiTT16^CSM1^.

### KiTTs orthologous to conventional kinetochore components are highly divergent

To better understand why several KiTTs orthologous to conventional kinetochore components were missed in all previous deep homology searches, we assessed the degree of sequence divergence of the KiTTs compared to other orthologs from across the eukaryotic tree of life. To this end, we generated phylogenetic trees of each of the orthologous groups using the orthologous groups previously delineated by us [16], and manually rooted these trees on the LECA node in accordance with the topology by Williamson *et al* [66]. This approach allows for the quantification of branch lengths from the LECA node to extant orthologs (**Figure 4A**). Branch lengths are a direct representation of sequence divergence. Plotting branch lengths of identified orthologs as a frequency distribution showed that all *T. thermophila* kinetochore orthologs had particularly long branch lengths compared to other orthologs, showing indeed high levels of divergence (**Figure 4B**). These observations underscore the fact that these *T. thermophila* kinetochore components evaded previous detection due to their high levels of divergence, despite the use of sensitive homology detection tools. Furthermore, this underscores the high rates of evolution of kinetochore proteins across diverse eukaryotic lineages.

**Figure 4.**
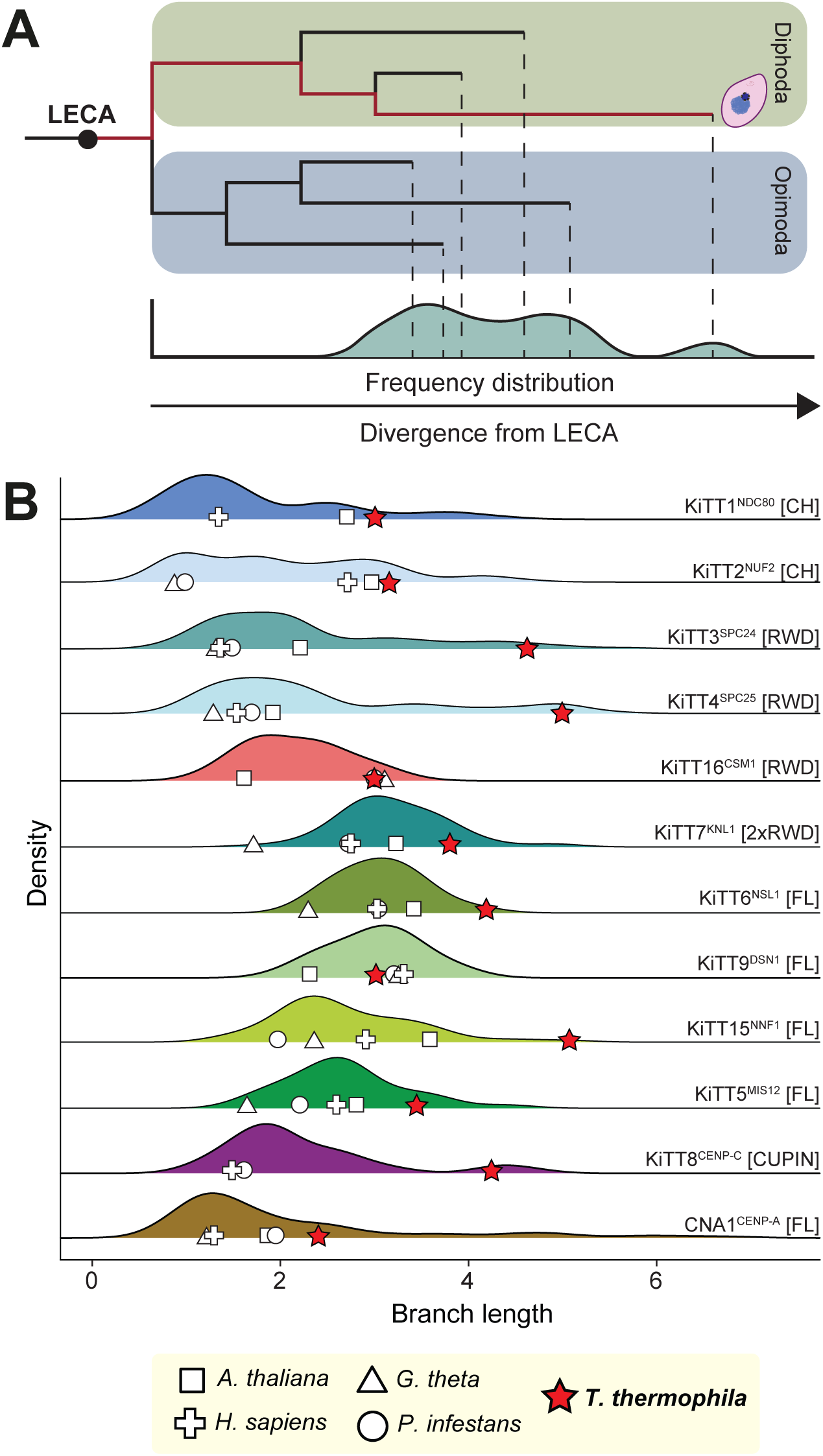
Sequence divergence analysis of T. thermophila kinetochore orthologs compared to other eukaryotes. (A) Schematic representation of a rooted phylogenetic tree and its corresponding branch length density plot. Gene trees are rooted at the inferred root of the eukaryotic tree of life according to Williamson et al, specifically, between Diphoda and Opimoda [66,123,124]. (B) Ridgeline plots of branch lengths of kinetochore orthologs samples across eukaryotic diversity. Phylogenetic trees are inferred based on full-length (FL) or domain-specific alignments (CH/RWD/CUPIN). Alignments were made by adding the KiTT sequence to a set of orthologs sampled across eukaryotes as defined by Tromer et al. [16]. Branch lengths are measured from the tips to the inferred root as described in (A). Branch lengths of orthologs from *T. thermophila* are indicated with a red star. For comparison, orthologs from five additional species are highlighted (see in-figure legend), representing divergent lineages of eukaryotes. *H. sapiens*: *Homo sapiens* (Metazoa), *A. thaliana*: *Arabidopsis thaliana* (Archaeplastida), *P. infestans*: *Phytophthora infestans* (Stramenopila;SAR), *G. theta*: *Guillardia theta* (Cryptista), *T. thermophila* (Alveolata;SAR).

### Positional architecture of the KiTTs in the T. thermophila kinetochore

To investigate the architecture of the kinetochore in *T. thermophila,* we next determined their locations in metaphase MICs relative to the inner kinetochore protein KiTT8^CENP-C^ and the outer kinetochore KiTT1^NDC80^, representing opposing ends of the kinetochore structure. We did so by performing super resolution immuno-imaging of the C-terminal HA/mCherry-tags on KiTTs in combination with the custom antibodies against the C-terminus of KiTT8^CENP-C^ or N-terminus of KiTT1^NDC80^ ^[67]^ (**Figure 5A-5B**). The mean distances of the N-terminus of KiTT1^NDC80^ to the C-termini of KiTT3^SPC24^ and KiTT4^SPC25^ were 53 ± 11 nm and 59 ± 18 nm (mean ± SD), respectively (**Figure 5C)**. This is consistent with the reported lengths of the NDC80 complexes of budding yeast (∼57 nm, [68]) and human (∼62 nm; [69]). Compared to the NDC80 complex, the C-termini of the MIS12 complex components KiTT5^MIS12^, KiTT6^NSL1^ and KiTT9^DSN1^, and KiTT7^KNL1^ placed slightly closer to the inner kinetochore, indicating a conformation similar to the human KMN network [4,56,70]. Therefore, the general architecture of the *T. thermophila* kinetochore closely resembles that of well-characterised opisthokont systems, validating our approach and showing that this archetypical conventional architecture was maintained despite high levels of sequence divergence.

**Figure 5.**
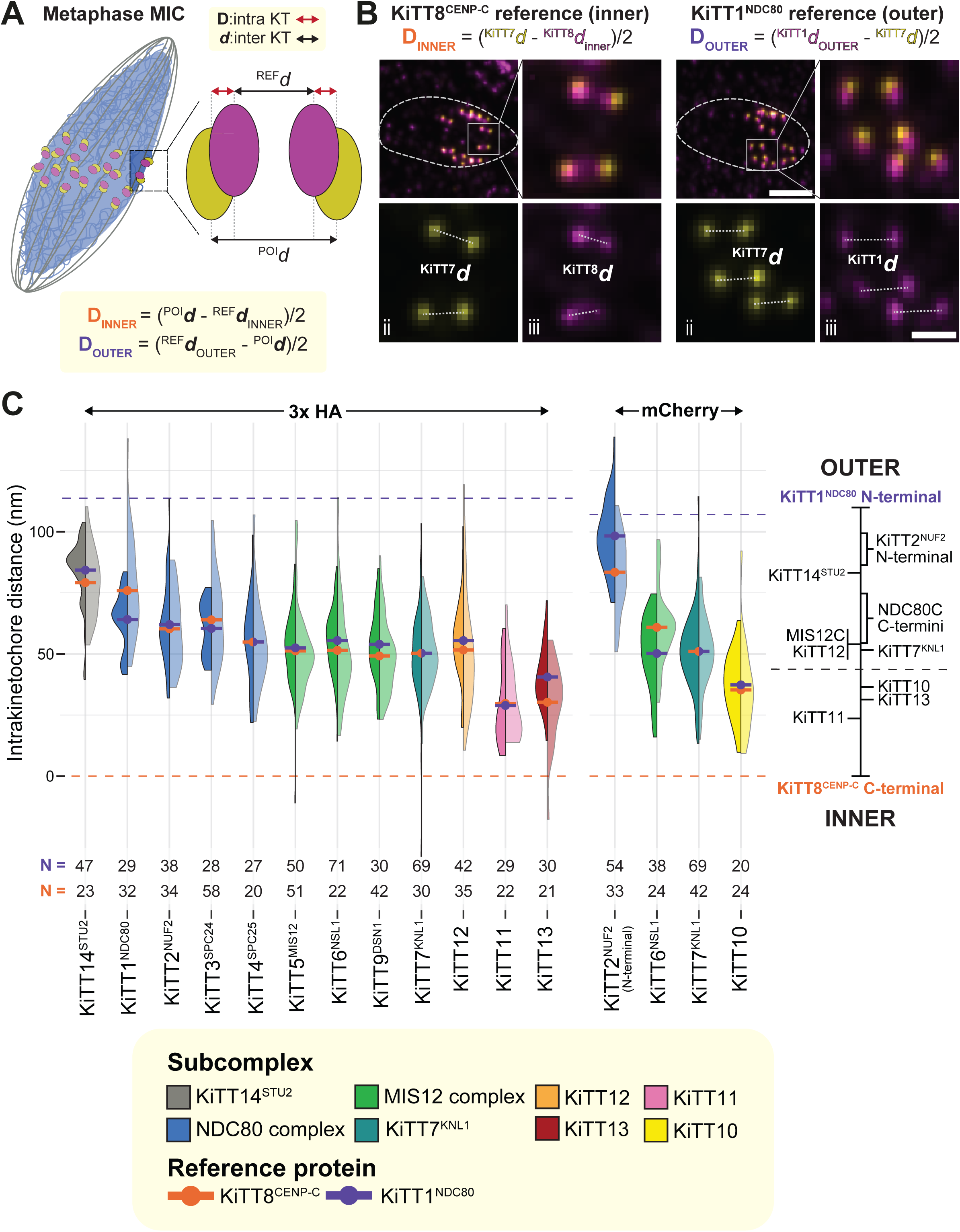
Spatial organization of KiTTs relative to KiTT8^CENP-C^ and KiTT1^NDC80^. (A) Schematic representation of kinetochores in a metaphase MIC. The inset shows a reconstruction of a kinetochore pair stained for a protein of interest (POI) and an inner kinetochore reference protein. To determine the intra-kinetochore distance (D), bioriented kinetochore pairs were manually selected. First, the inter-kinetochore distance (d) is defined as the measurement between the centroids of the protein of interest and a reference protein, where KiTT1^NDC80^ is the reference for the outer kinetochore, and KiTT8^CENP-C^ as the inner kinetochore reference. The intra-kinetochore distance is then calculated by subtracting the inter-kinetochore distance of the reference protein from that of the protein of interest, or *vice versa*, for the measurements taken using the outer reference protein or the inner reference protein, respectively, and is then divided by two. (B) Representative images of cells with 3xHA tags on KiTT7^KNL1^ (yellow) co-stained with custom antibodies for KiTT8^CENP-C^ (magenta) or KiTT1^NDC80^ (magenta). Insets i, ii, iii show zoom-ins on kinetochore pairs. White dotted lines in insets i, ii and iii represent the measured distances resulting in dKiTT7^KNL1^ and dKiTT1^NDC80^ (right)/dKiTT8^CENP-C^ (left). The white dotted line in the top-left images depict the outline of the MIC. Scale bar image top-left 2 μm, zoom-ins 0.5 μm. (C) Plot showing the mean distances of all KiTTs relative to KiTT8^CENP-C^ and KiTT1^NDC80^. The bars represent the mean distances measured from either the KiTT1^NDC80^ reference (purple), or the KiTT8^CENP-C^ reference (orange). The violin plots depict the distribution of the individual measurements obtained from the KiTT1^NDC80^ reference (left, darker shade), or the KiTT8^CENP-C^ reference (right, lighter shade). On the right side of the plot, a rough outline of the average position of each KiTT is placed in the general architecture of the *T. thermohila* kinetochore. The horizontal dotted lines represent the KiTT8^CENP-C^ reference point (below, at 0), or the sum of the distance from the KiTT8^CENP-C^ reference point to KiTT7^KNL1^ and KiTT7^KNL1^ to KiTT1^NDC80^ (above). This latter point was taken to rescale the KiTT1^NDC80^-based measurements, effectively representing the outer limit of the kinetochore based on our measurements. The numbers on the top and bottom of the plot refer to the number of measured pairs across the KiTT1^NDC80^- and KiTT8^CENP-C^-based analyses, respectively.

We next examined the relative position of the unconventional KiTTs. The C-terminus of KiTT12 localised at the outer kinetochore at ∼55 nm from KiTT8^CENP-C^, near the MIS12 complex. Conversely, KiTT10, KiTT11 and KiTT13 showed a more centromere-proximal localisation (∼37nm, ∼29nm, ∼41nm from KiTT8^CENP-C^, respectively) consistent with an inner kinetochore localisation of these proteins in *T. thermophila* (**Figure 5C**).

### Diverse origins of the unconventional inner kinetochore components KiTT10, 11 and 13

To investigate the evolutionary origin and putative function of the unconventional KiTTs 10/11/13 at the inner kinetochore, we performed broad deep homology searches across prokaryotic and eukaryotic proteomes coupled with phylogenetics. HMM-v-HMM searches of KiTT10 (TTHERM_00056030) revealed homology to WD40 domain-containing proteins, which was verified by structure predictions for KiTT10 using ESMfold [71] (**Figure 6A**). Additional searches using both a profile HMM based on KiTT10 orthologs in the 10 *Tetrahymena* species and the predicted AF2 structure of KiTT10 allowed for the identification of a putative ortholog in the ciliate *Paramecium tetraurelia*, which lacks the large insertions seen in *T. thermophila* KiTT10 (**Figure 6A**). In eukaryotes, the WD40 domain is one of the most abundant domains in proteins at the heart of many different cellular processes [72]. Due to the extensive divergence of both the *T. thermophila* and *P. tetraurelia* KiTT10 orthologs, we were unable to further specify to which eukaryotic WD40 subfamily KiTT10 belongs.

**Figure 6.**
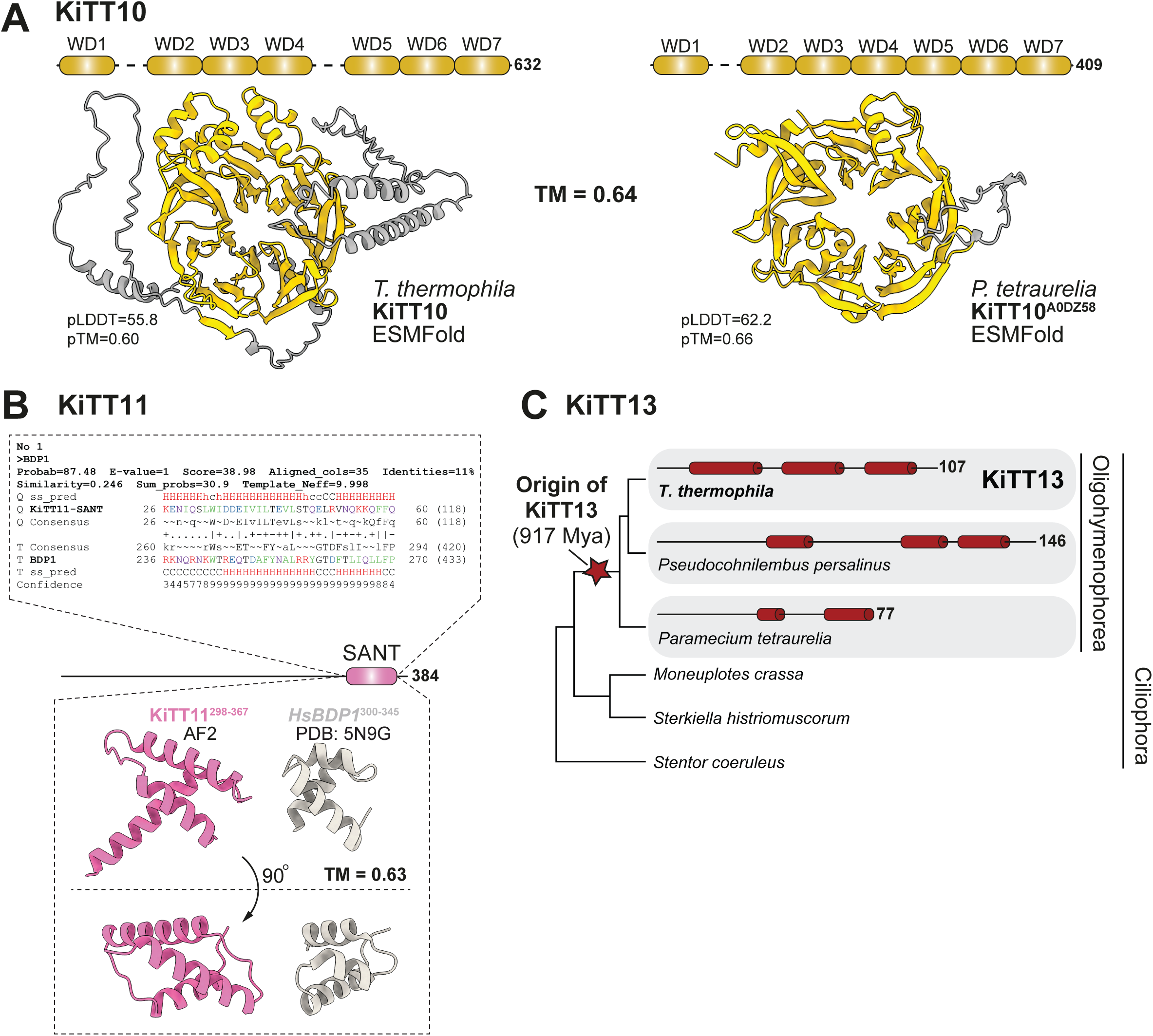
Diverse origins of unconventional inner kinetochore components in T. thermophila. (A) ESMFold prediction of KiTT10 and its putative *P. tetraurelia* ortholog with TM-score of their alignment (middle). (B) Top: HHsearch alignment of the KiTT11 SANT domain model to a eukaryote-wide BDP1 profile. Bottom: AF2 prediction of the KiTT11 SANT domain shown side-by-side to the SANT domain of the human BDP1 ortholog (PDB ID: 5N9G[125]). (C) Schematic representation of KiTT13 orthologs and their secondary structures across Ciliophora. Red cylinders represent α-helices and the number represents the total length of the amino acid sequences of the protein. The diagrams are drawn to scale. Absence of secondary structure diagrams indicate the absence of a KiTT13 ortholog. The red star indicates the inferred origin of KiTT13 at the base of oligohymenophorea [73].

Using AF2, we predict KiTT11 (TTHERM_00463460) to have a helix-turn-helix domain in the C-terminal region (**Figure 6B**). Sequence-based homology searches verified this and identified the SANT domain of Bpd1, a component of the TFIIIB transcription initiation complex, as having the strongest similarity to the KiTT11 helix-turn-helix. Subsequent phylogenetic analysis revealed that Bdp1 duplicated repeatedly in ciliates, particularly among Oligohymenophorea, where we infer one *Tetrahymena*-specific duplication led to the origin of KiTT11, while Bdp1 is typically present as single-copy orthologs in Stramenopila and Myzozoa (**Figure S7**). From this, we concluded that there are no 1-to-1 orthologs of KiTT11 outside of the *Tetrahymena* genus, and it is likely that it acquired its kinetochore localisation following this duplication. Therefore, we hypothesise that KiTT11 acquired its kinetochore function via neofunctionalization in the midst of an expansion of the Bdp1 gene family in ciliates, and pinpoint it to a *Tetrahymena*-specific duplication (**Figure S7**).

Homology searches for KiTT13 (TTHERM_00218939) found clear orthologs in *P. tetraurelia* and *Pseudocohnilembus persalinus*, which are both member of the ciliate class Oligohymenophorea, to which also *T. thermophila* belongs. Curiously, outside of Oligohymenophorea, we could not find homology to any other protein in a broad set of eukaryotic proteomes, including in other ciliates (**Figure 6C**). Based on this highly limited phylogenetic distribution, as well as its short length (107 residues in *T. thermophila*) and lack of complex folded domains, we inferred that KiTT13 is a *de novo* gene that arose in the Oligohymenophorea lineage, at least 827 Mya [73], followed by functionalization into the *T. thermophila* kinetochore (**Figure 6C**).

Combined, we find that the unconventional components that complement KiTT8^CENP-C^ at the inner kinetochore of *T. thermophila* have disparate evolutionary origins. Yet, they likely co-orchestrate the various functions of the inner kinetochore of *T. thermophila*.

### The outer kinetochore protein KiTT12 is a kinesin required for accurate chromosome segregation and NDC80 residency at kinetochores

Our deep homology searches showed that the only unconventional KiTT that resides at the outer kinetochore, KiTT12, contains a highly-divergent N-terminal kinesin motor domain (**Figure 7A**). To trace its evolutionary origin, we aligned the sequence of its inferred motor domain to the set of kinesin motor domains of Wickstead et al [74] and generated a phylogenetic tree to place KiTT12 in its larger kinesin family (**Figure 7A**). In addition, we used AF2 to predict motor domain structure of all these kinesins and generated a structure-based phylogeny (**Figure S8**). Both phylogenies placed KiTT12 within the kinesin-6 clade. Kinesin-6 family kinesins have roles in cytokinesis during mitosis in opisthokonts and Amoebozoa, but no role at the kinetochore has been reported in these lineages [75–78].

**Figure 7.**
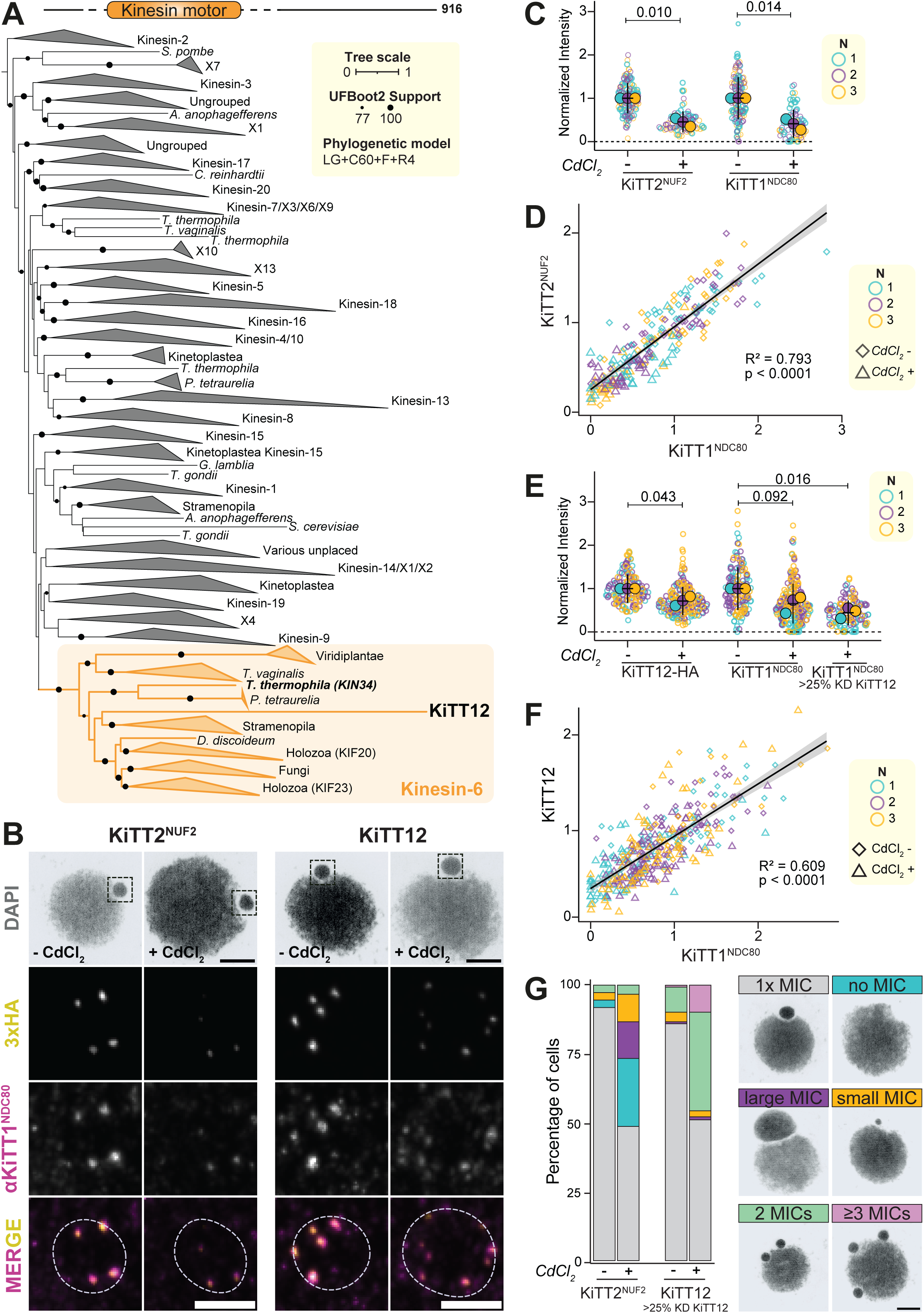
KiTT12 is a kinesin of the Kinesin-6 family and is important for KiTT1^NDC80^ occupancy. (A) Phylogenetic tree of the motor domain of KiTT12 with the motor-domain alignment of Wickstead et al. [74]. The tree was inferred using the LG+C60+F+R4 model. KiTT12 is not the only kinesin-6 family member in *T. thermophila*. KIN34 (TTHERM_00637750), also part of the proximity proteome of the kinetochore (Figure 2), is placed within this clade in our phylogenies. (B) Immunofluorescence staining of KiTT2^NUF2^ (left) and KiTT12 (right). RNAi was induced by addition of CdCl_2_ into the culture 48 h prior to the fixation. The cells were stained for KiTT2^NUF2^-HA/KiTT12-HA (yellow) and KiTT1^NDC80^ (magenta) and DAPI (gray). Insets show magnified micronuclei. All images are maximum intensity projections. Scale bars, 5 µm, MIC zoom-ins, 2 µm. (C) Quantification of KiTT2^NUF2^ and KiTT1^Ndc80^ immunostaining intensity in control cells (-CdCl_2_) and RNAi-induced cells (+ CdCl_2_). RNAi expression was induced by addition of CdCl_2_ into the culture 48 h prior to the fixation. Data was obtained in three independent experiments, as specified in the in-figure legend. (D) Scatter plot with fitted linear model showing the correlation between intensities of KiTT2^NUF2^and KiTT1^Ndc80^ signal from the immunostained images. Each dot represents a single MIC. Both RNAi-induced and RNAi-uninduced cells are shown across three independent replicates, as specified in the in-figure legend. (E) Same as (**C**) but for KiTT12 RNAi. Right-most data on KiTT1^Ndc80^ intensity represents only cells with >25% knockdown of KiTT12. (F) Same as (**D**) but for KiTT12 RNAi. (G) Stacked barplot showing the prevalence of various phenotypic effects of knockdown of KiTT2^NUF2^ (left) and KiTT12 (right) in cells where RNAi is not induced (-CdCl_2_) versus induced (+ CdCl_2_) as specified in in-figure legend. The bars are color-coded according to the phenotypes as shown in the in-figure legend. Scale bar in-figure legend: 5 µm. Statistical significance was determined by Student’s t-test in **C** and **E**, and by linear modelling in **D** and **F**.

To investigate the functions of the unconventional *T. thermophila* kinetochore components, we used CdCl_2_-inducible RNA interference (RNAi)-mediated knockdown [79]. We obtained significant knockdown of KiTT12 and KiTT2^NUF2^ (**Figure 7B**), but not KiTT10, 11 and 13. As expected [80], RNAi of KiTT2^NUF2^ caused a significant reduction in kinetochore levels of KiTT1^NDC80^ (p = 0.014, **Figure 7C**), and the levels of KiTT2^NUF2^ significantly correlated with those of KiTT1^NDC80^ at kinetochores across MICs (p <0.001, **Figure 7D**). Interestingly, we found that depletion of KiTT12 (>25%) also resulted in a significant reduction of KiTT1^NDC80^ levels at kinetochores (p = 0.016, **Figure 7E**). Accordingly, and similar to KiTT2^NUF2^, KiTT12 levels at kinetochores significantly correlated with those of KiTT1^NDC80^ (p<0.0001, **Figure 7F**). A subsequent assessment of MIC morphology in KiTT2^NUF2^ RNAi cells showed that the reduced levels of KiTT1^NDC80^/KiTT2^NUF2^ caused substantial MIC aberrations or complete MIC absences (**Figure 7G**), indicative of chromosome segregation errors and consistent with the phenotypes upon CNA1 knock-downs [35,36]. The MICs in KiTT12 RNAi cells similarly showed substantial aberrations (**Figure 7G**). These aberrations were different from those in KiTT2^NUF2^ RNAi cells, however, suggesting potentially additional functions for KiTT12, such as a role in karyokinesis/cytokinesis as previously described for kinesin-6 family members.

## DISCUSSION

> “It turns out that above and beyond the immense morphological and ecological diversity of ciliates, their most bizarre features are those hidden in the molecular realm.” (Boscaro & Keeling) [33]

We have used a combination of proximity proteomics, super-resolution imaging and deep homology detection to uncover the evolutionary hybrid kinetochore of the model ciliate *T. thermophila.* We show that its kinetochore is composed of at least three highly-conserved, nine cryptic conventional and four unconventional components (**Figure 8**).

**Figure 8.**
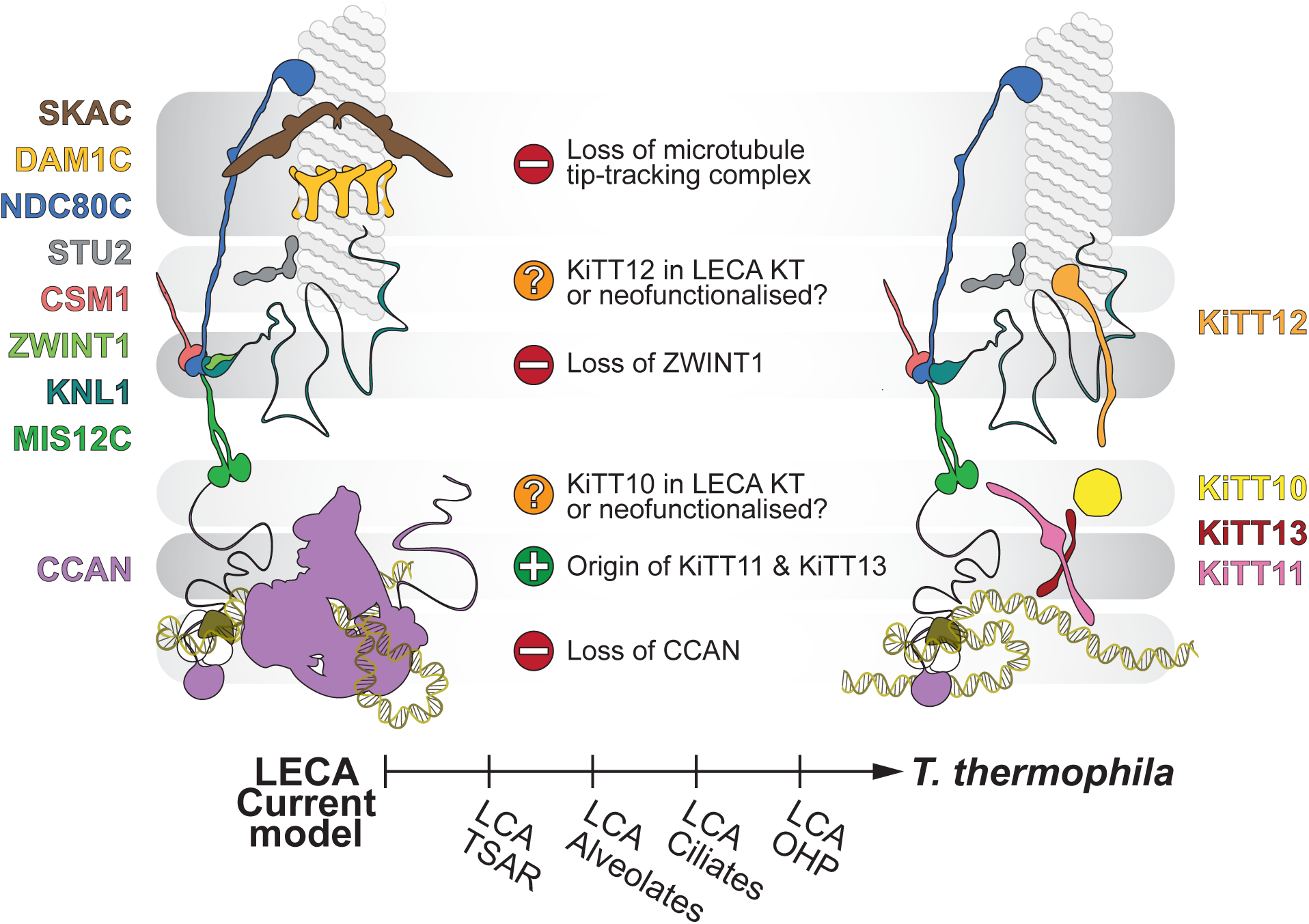
Model of the LECA kinetochore compared to our inferred T. thermophila kinetochore. Comparison between the inferred composition of the LECA kinetochore and the *T. thermophila* kinetochore. Major evolutionary transitions are described between the two kinetochore models. CSM1 is now inferred to be present in the LECA kinetochore based on its presence in the *T. thermophila* kinetochore as well as that in several other species (see discussion). Below describes the evolutionary lineage leading from LECA up to *T. thermophila*. Each transition occurred somewhere along this trajectory spanning ∼1.5-2 Bya. Locations of the components in the *T. thermophila* kinetochore model are based on the measurements from our intra-kinetochore measurements (Figure 5). LCA: Last common ancestor; TSAR: Telonemia, Stramenopila, Alveolata, Rhizaria; OHP: Oligohymenophorea.

In contrast to the previously-reported pervasive loss of kinetochore complexity in *T. thermophila* [15], we find that the bulk of its kinetochore consists of a highly-divergent, yet near-complete, KMN network (9 subunits out of 10, with the exception of ZWINT1) and a single CCAN component KiTT8^CENP-C^. Strikingly, these components contain most domains and motifs that are required for the molecular interactions of the KMN-CENP-C axis as found in established models like human and yeast, suggesting they have maintained their conventional kinetochore functions despite extreme divergence of their amino acid sequence. This architecture, including loss of most CCAN subunits following divergence from LECA [15], is quite pervasive across the eukaryotic tree of life, as it is found in evolutionary-distant model organisms like *D. melanogaster* (fly) [18]*, C. elegans* (worm) [81], *A. thaliana* (plant) [82], and of the alveolate sister clade to ciliates, the apicomplexans (i.e. *Plasmodium* and *Toxoplasma*) [28,29]. The latter species also have a highly-reduced inner kinetochore, extremely-divergent KMN (KNL1-MIS12-NDC80) network subunits, and also contain lineage-specific duplicates of CENP-C [28]. The conserved axis of the *T. thermophila* kinetochore is therefore exemplary of recurrent convergent evolution of such a kinetochore composition across diverse eukaryotic lineages.

Beyond this conserved axis, the *T. thermophila* kinetochore has numerous seemingly non-canonical features, further exemplifying compositional plasticity of the kinetochore across eukaryotic evolution. For example, we found no subunits of the DAM1 or SKA complexes, suggesting a striking absence of accessory microtubule tip-tracking complexes. Possibly, their role in stabilising outer-kinetochore attachment to microtubules is fulfilled by the kinesin-6 family member KiTT12, which likely binds microtubules through its motor domain and is important for NDC80 levels at the outer kinetochore. To our knowledge, ours is the first description of a role for a kinesin-6 family member in kinetochore function. Kinesin-6 proteins are more commonly associated with roles in cytokinesis, from studies using slime molds and human cells [75–78]. These species are more closely-related to each other, grouped together in the so-called Amorphea, than they are to *T. thermophila [14]*. Combined, this implies that the localisation and function of KiTT12 at the *T. thermophila* outer kinetochore is either a derived feature specific to the lineage leading up to *T. thermophila*, or equally possibly, that this reflects an ancestral function of the kinesin-6 family, which might have been specifically lost in Amorphea.

Our BioID experiments reveal the presence of several components of the y-TuRC complex in close proximity to kinetochores in *T. thermophila*. It is possible that this pool of y-TuRC localised near kinetochores in *T. thermophila* has a role in chromosome congression and microtubule nucleation, in striking parallel to a recently-described mechanism in mammalian cells [83]. Additionally, we find that KiTT14^STU2^, an ortholog of STU2/CKAP5/ch-TOG/XMAP-215, localises to the outer kinetochore in *T. thermophila*. STU2 is a plus-end microtubule-associated protein with a microtubule polymerase function, regulating microtubule dynamics at multiple locations in the cell [84], including a prominent role at the y-TuRC complex [85]. A role for STU2/ch-TOG has recently been reported at the kinetochore in Metazoa [86], and its apparent kinetochore localisation *Plasmodium berghei* [28] and budding yeast [87] make it likely that STU2 is an ancestral kinetochore component tracing back to LECA.

Surprisingly, we identified a *bona fide* ortholog of the yeast monopolin subunit CSM1 in our BioID experiments, showing that this protein associates with kinetochores in our exclusively-mitotic *T. thermophila* cultures. Indeed, previous work on the orthologous proteins Pcs-1 (fission yeast) [88], CENP-68 (slime mold) [89], and more recently TITAN-9 (plants) [90] and AKiT4 (*P. berghei) [28]* now strongly suggest that CSM1 orthologs are part of mitotic as well as meiotic kinetochores and that this protein has to be considered a *bona fide* eukaryotic-wide kinetochore protein that was part of LECA, confirming our previous bioinformatic predictions [91]. Although at the kinetochore, the absence of a monopolin-binding motif in the N-terminal disordered tail of the DSN1 ortholog KiTT9, indicates it is most likely differently recruited to kinetochores compared to other eukaryotes [91]

The inner kinetochore is responsible for organising centromeric chromatin as well as connecting centromeric chromatin to the outer kinetochore, and has been recognised as a hub for evolutionary novelty and is repeatedly remodelled across eukaryotes [92]. For instance, the inner kinetochore of the silk moth *Bombyx mori* is rewired [93], the inner kinetochore of the yeast *C. neoformans* is replaced with a novel protein called bridgin [20], and lineage-specific components that interact directly with the centromeric DNA are found in mammals (CENP-B) [94] and budding yeast (CBF3 complex) [95]; the latter which has recently been shown to even dictate the evolutionary dynamics of centromeric DNA in yeast [96]. Consistent with these patterns, the *T. thermophila* inner kinetochore is enriched for evolutionary novelties: KiTT11 and KiTT13 derive from a *Tetrahymena*-specific duplication and an Oligohymenophorea-specific gene-birth event, respectively, while the WD40 protein KiTT10 may represent either an extremely-divergent member of an ancestral centromere-proximal factor family (e.g. BUB3 or RBBP7 [97]), or an Oligohymenophorea-specific innovation. Alternatively, KiTT10 could represent a member of a centromere-proximal factor that was present in LECA, but is absent from well-studied model organisms. However, as noted, we have not been able to find homologs of KiTT10 in other species, preventing the exploration of these options. Notably, KiTT11, KiTT13 and the conserved CENP-A loading factor MIS18 all show similar G2/M-enriched expression profiles (**Figure 2E**), and KiTT11 contains a predicted SANT-like domain reminiscent of those found in CENP-A loading modules such as KNL2/MIS18BP1. Screening for putative interacting KiTTs using AF2, the co-fold of KiTT11 and KiTT13 had a reasonable interface predicted template modelling score suggesting that these two lineage-specific components may form a novel complex at the inner kinetochore (iPTM: 0.619; **Figure S9**). However, we note that KiTT13 emerged prior to the duplication that led to the origin of KiTT11 (**Figure 6C, Figure S7**), suggesting that KiTT13 may also have a KiTT11-independent function. Together, these observations raise the possibility that these *Tetrahymena*-specific components act alongside MIS18 in centromeric chromatin maintenance or CENP-A deposition, underscoring the distinctive evolutionary innovations present at the *T. thermophila* inner kinetochore.

The presence of conventional kinetochore subunits in *T. thermophila* that went unnoticed in earlier investigations is revealing, both on the limitations of such analyses on larger evolutionary scales (i.e. eukaryote-wide characterisation, compared to our focus on a single species), as well as methodological advances in the detection of highly-divergent orthologs and the availability of a large amount of predicted proteomes. Importantly, we employed highly-sensitive HMM-v-HMM searches and targeted these to a set of experimentally-determined kinetochore candidates, which we obtained through the pioneering use of a BioID approach in *Tetrahymena*. Furthermore, newly-developed protein structure prediction methodologies such as AF2, AF3 and ESMFold [71,98,99] proved highly valuable in assessing putative orthologs and discovering distant homologies. We also note the fact that the predicted proteome of *T. thermophila* has undergone several version updates over the last decade, with notably the most recent reannotation of the *T. thermophila* genome reporting close to 10% of newly-predicted genes that were missed from the 2021 version, including the orthologs of CSM1 and NNF1 that we had predicted based on our homology searches [60]. Such missing predictions of existing genes are an important consideration when inferring absences of proteins from species [100], and our observations here highlight the significance of a careful examination of inferred absences in original genome assemblies. Crucially, both the sequence-based HMM-v-HMM as well as the structure-based homology-detection methods we employed relied in large part on the availability of a large set of predicted proteomes from multiple *Tetrahymena* species, underscoring the importance of deep taxon-sampling for comparative genomics studies and sensitive homology assessment.

## OUTLOOK

Our findings show that the *T. thermophila* kinetochore—previously inferred to be highly reduced—is instead built around a largely conventional KMN–CENP-C axis composed of deeply divergent orthologs supplemented with only a small number of lineage-specific factors. This reveals that ciliates, despite their high levels of divergence and distinctive nuclear biology, adhere more closely to ancestral kinetochore architecture than anticipated. Rather than representing an exceptional departure from canonical machinery, *T. thermophila* exemplifies a broader evolutionary principle: LECA-derived protein complexes are generally retained across eukaryotes, even when their subunits undergo extreme divergence or acquire lineage-specific innovations. A similar pattern has been observed for the ciliate Mediator complex, which preserves conserved core subunits but incorporates *Tetrahymena*-specific components to possibly accommodate its unusual nuclear context [101,102].

Methodologically, this work highlights the power of combining proximity proteomics with lineage-focused homology detection and structure prediction in non-model systems. Many of the cryptic orthologs identified here would have remained undetected by sequence-based homology searching approaches alone, underscoring the need for experimental anchoring when interpreting inferred absences in divergent genomes. Further expansion of the genetic and imaging toolkit in *Tetrahymena*—including more versatile conditional perturbation strategies and live-cell micronuclear imaging—will be essential to dissect the functions of its unconventional KiTTs (e.g., KiTT10-11-13, KiTT12) and to illuminate how deeply conserved cellular systems are remodelled over vast evolutionary timescales.

## LIMITATIONS OF THIS STUDY

Working with *T. thermophila* imposes methodological constraints typical of protist models, which are less developed than more standard model organisms. While we expanded its genetic toolbox by establishing TurboID, new tagging constructs, and optimized micronuclear isolations for BioID, several approaches remain limited compared to animal and fungal systems. Functional interrogation of the unconventional KiTTs—such as precise mechanistic roles of KiTT10/11/13 or the kinesin-6 KiTT12—is constrained by the limited efficiency of inducible knockdowns, the lack of robust degron systems, and the difficulty of live-cell imaging of micronuclear kinetochores at high temporal resolution. Likewise, evolutionary inferences and compositional reconstructions are partly shaped by genome annotation quality, which continues to improve but remains incomplete. These limitations underscore that deeper mechanistic insight will require continued development of experimental methods and proper genomic annotation in ciliates and other non-model eukaryotes.

## MATERIALS AND METHODS

### Cell Culture

*Tetrahymena thermophila* strains CU427.4 (TSC_SD00715) and B2086.2 (TSC_SD00709) were obtained from the *Tetrahymena* stock center (https://tetrahymena.vet.cornell.edu). Cells were cultured in super proteose peptone (SPP) medium (2% proteose peptone, 0.1% yeast extract, 0.2% glucose, 0.003% Fe-EDTA) supplemented with 100 µg/ml Normocin (InvivoGen, Cat# ant-nr-05) at 30 °C.

### Protein tagging

For the endogenous tagging of KiTT1^NDC80^ with 3xHA-TurboID, the EGFP tag and KiTT1^NDC80^ fragment of KiTT1^NDC80^-EGFP-NEO4 construct (a gift from Dr. Rachel Howard-Till [44]) were removed with NotI/SpeI and replaced with PCR amplified KiTT1^NDC80^ and 3xHA-TurboID amplified from a codon-optimized 3xHA-TurboID gBlock (IDT). The C-terminal 3xHA-TurboID constructs for all other genes were generated as previously described [103]. In short, sequences corresponding to the C-terminal end of the genes and downstream UTRs were amplified from genomic DNA of wild type cells with the primers listed in Table S4. These fragments were integrated to flank the C-terminal tagging module of 3xHA-TurboID-NEO4. To create KiTT2^NUF2^-3xHA-TurboID construct, PCR fragments of the C-terminal end and downstream UTR of KiTT2^NUF2^ were cloned into pHA-NEO4 [103] using BamHI/SacI and KpnI/HindIII restriction sites. HA tag was excised with BamHI/SpeI and replaced with 3xHA-TurboID. The sequences of KiTT2^NUF2^ and UTR of KiTT2^NUF2^-3xHA-TurboID were removed with BamHI/SacI or EcoRV/KpnI and replaced with PCR amplified fragments of KiTT5^MIS12^, KiTT11, KiTT12. To create 3xHA-TurboID fused KiTT6^NSLl1^, KiTT7^KNL1^, KiTT9^DSN1^, KiTT10, KiTT13, KiTT14^STU2^, TTHERM_00932010, the gene and UTR sequences of pKiTT11-3xHA-TurboID were digested out with BamHI/SacI and EcoRV/KpnI and replaced with PCR amplified sequences of the corresponding genes. For KiTT3^SPC24^-3xHA-TurboID and KiTT4^SPC25^-3xHA-TurboID constructs, the 3xHA-TurboID, gene and UTR fragments were combined by overlapping PCR. To create KiTT2^NUF2^-3xHASpot, the KiTT2^Nuf2^-3xHA sequence was amplified from KiTT2^Nuf2^-3xHA-TurboID construct with a primer set including Spot. The tag of KiTT2^Nuf2^-HA construct was removed with SacI/SpeI and replaced with the KiTT2^NUF2^-3xHASpot fragment. For the C-terminal mCherry tagged proteins, the 3xHA-TurboID sequence of KiTT6^NSL1^, KiTT7^KNL1^, KiTT10, was removed with BamHI/PstI and replaced with mCherry sequence from pmCherry-NEO4 [103]. To create N-terminally tagged mCherry-KiTT2^NUF2^, the N-terminal and upstream UTR sequences were amplified from the genomic DNA of wild type cells. These fragments were integrated into pBNMB-mNeonGreen (a gift of Dr. Miao Tian) at BamHI/KpnI and SacI/SalI sites respectively. The mNeonGreen tag was excised with NdeI/KpnI and replaced with PCR amplified N-terminal sequence of KiTT2^NUF2^ and mCherry tag using Gibson assembly. To generate N-terminally tagged mNeonGreen-TTHERM_00393260^MADBUB^, the MTT1-mNeonGreen-NEO5 cassette was digested with BamHI/SalI from pBNMB-mNeonGreen and combined with PCR amplified regions of N-terminal end and upstream 5’UTR using overlapping PCR. All constructs were transformed into the wild type cells by biolistic transformation as previously described [104].

### RNAi knockdown

For KiTT2^NUF2^ and KiTT12 RNAi constructs, a target sequence of the gene of interest was amplified from wild type genomic DNA with two sets of primers listed in **Table S4**. These fragments were cloned into a hairpin expression cassette that contains a CdCl_2_-inducible MTT1 metallothionein promoter for hairpin expression and cycloheximide resistance [105,106]. The RNAi constructs of KiTT2^NUF2^ and KiTT12 were introduced into the cells expressing KiTT2^NUF2^-3xHASpot and KiTT12-3xHA-TurboID respectively. Cells were selected with increasing concentration of cycloheximide. For RNAi experiments, cells were treated with 1 µg/ml CdCl_2_ for 48 h to induce hairpin expression.

### Phylogenetic analyses

Phylogenetic analyses were performed by aligning homologous sequences using MAFFT (v.7.520, E-INS-i/L-INS-i) [107]. Alignments were trimmed using trimAl v1.4.rev15 [108] and sequences with low occupancy were filtered out using a custom Python script. Based on the obtained MSAs, maximum likelihood phylogenies were inferred using IQ-TREE v2.3.6 [109]. Structure-based phylogenetic analysis was performed by structure prediction using AF2 [98,110]. A phylogenetic tree was inferred based on the predicted structures using FoldTree [111].

### Divergence analysis

Kinetochore orthologs representing the breath of eukaryotic diversity were obtained from Van Hooff *et al.* 2017 [15]. The corresponding *T. thermophila* ortholog was added to the eukaryote-wide set and sequences were aligned using MAFFT (v.7.520, L-INS-i) [107]. MSAs were then trimmed using trimaAl with -gt 0.5 and all sequences with <50% occupancy over the length of the MSA were removed using a custom Python script. Then, maximum likelihood phylogenies were inferred using IQ-TREE v2.3.6 [109]. The obtained phylogenetic trees were manually rooted in accordance to Williamson et al 2024 [66], with the caveat that single-protein alignments rarely suffice to produce faithful species phylogenies. However, these erroneous topologies have only marginal effects on branch length inferences. Finally, branch lengths were quantified using the ete3package [112].

### HMM-vs-HMM homology searches

OrthoFinder (Emms & Kelly, 2019) (v.2.5.5, -S blast; otherwise default settings) was used to generate OGs from the predicted proteomes of 10 *Tetrahymena* species [38]. OG sequences were then aligned using MAFFT (v.7.520, E-INS-i) [107]. The alignments were then converted into an HHsuite-type database of HMM profiles using the HHsuite3 toolkit [113]. First, consensus sequences were predicted using the hhconsensus method. Then, alignments were converted into a3m format and predicted secondary structure was added using the HHsuite3 Perl scripts reformat.pl and addss.pl, respectively (downloaded from https://github.com/soedinglab/hh-suite/tree/master/scripts, last accessed March 2nd 2024). Finally, profile-vs-profile HMM searches (hhsearch) were performed using the *Tetrahymena* profiles as a query against a curated kinetochore database [22] supplemented with the KOG/COG database (downloaded from http://ftp.tuebingen.mpg.de/pub/ebio/protevo/toolkit/databases/hhsuite_dbs/, last accessed March 22nd 2024).

### Protein structure prediction

Predicted protein structures were generated using AlphaFold2 (AF2) (Jumper et al., 2021), AlphaFold3 (AF3) [99], and ESMfold [71]. For AF2, custom MSAs were supplied, which has been shown to improve predictions of highly-divergent and/or poorly-sampled lineages [114]. Specifically, the alignments used as a query for the HMM-vs-HMM searches were used, consisting of a set of OrthoFinder-derived orthologs across the 10 Tetrahymena species (see HMM-vs-HMM homology searches). AF2 was run using the ColabFold implementation using default settings [98,110]. AF3 was run from the online implementation via https://alphafoldserver.com/. ESMfold was run through the ColabFold implementation using default settings [71,110]. Structures were visualised with ChimeraX v1.5 [115].

### Micronuclear isolation and TurboID

Micronuclei were isolated with a method adapted from [47]. Briefly, cells expressing 3xHA-TurboID tagged bait proteins and wild type cells were treated with 50 µM Biotin (Sigma-Aldrich, Cat# B4501) for 1 h at 30 °C and washed twice with 10 mM Tris-HCl pH 7.5. Cells were resuspended in Lysis buffer (0.1 M Sucrose, 2 mM MgCl_2_, 4% Gum arabic, 10 mM Tris, 1 mM Iodoacetamide, 10 mM Butyric acid, pH 6.75) supplemented with Complete Protease Inhibitor (Roche, Cat# 5056489001). Cells suspension was combined with 0.5 mm glass beads (BioSpec, 11079105) in a cold chamber and blended with BeadBeater (BioSpec) for 6x (30 sec ON and 30 sec OFF) at 4 °C. The lysate was combined with 0.63% Octanol (Sigma-Aldrich, Cat# 8209310100) and blended for 20 sec, followed by centrifugation at 5830 rpm at 4 °C for 28 min in a swinging-bucket rotor (JS-24.38 rotor, Beckman Coulter Avanti J-30I high-speed centrifuge). The pellet was resuspended in the Nucleus Wash Buffer (0.25 M Sucrose, 1 mM MgCl_2_, 3 mM CaCl_2_, 10 mM Tris, 1 mM Iodoacetamide, 10 mM Butyric acid, pH 7.5). The supernatant and pellicle layer containing MIC-MAC nuclei were transferred into the cold chamber with glass beads, blended for 20 sec and centrifuged at 5830 rpm for 28 min. After two rounds of blending - centrifugation at 5830 rpm for 28 min, pellets containing MICs were combined.

Nuclei were lysed in RIPA buffer (10 mM Tris-HCl pH 7.5, 150 mM NaCl, 0.5 mM EDTA, 0.1 % SDS, 1 % TritonX-100, 1 % NaDeoxycholate) supplemented with Complete Protease Inhibitor for 15 min on ice, sonicated 10x high power 15 sec ON and 45 sec OFF (Bioruptor Sonicator), and incubated with benzonase (Santa Cruz) for one hour at 4 °C. Lysates were clarified by centrifugation at 12 000 x g for 10 min at 4 °C. Protein concentration was determined with Bradford assay. Equal amount of supernatant was combined with Pierce streptavidin magnetic beads (Thermo Scientific, Cat# 88816) and incubated for 2 hours or overnight at 4 °C. Beads were washed once with 10 mM Tris-HCl pH 7.5, 150 mM NaCl, 0.5 mM EDTA, twice with 10 mM Tris-HCl pH 7.5, 500 mM NaCl, 0.5 mM EDTA, and twice with PBS.

For immunoblotting, proteins from 10 % of the beads were eluted with 2x SDS sample buffer, resolved by 10% SDS-PAGE gel and transferred onto nitrocellulose membrane. To detect the tagged proteins, the membrane was incubated with mouse/rabbit HA antibody (1:1000) and goat anti-mouse IgG (H+L)-HRP/goat anti-rabbit IgG (H+L)-HRP (1:5000, Bio-Rad, Cat# 1706516/1:5000, Bio-Rad Cat# 1706515). HRP-conjugated Streptavidin (1:1000, Abcam, Cat# ab64269) was used to detect biotinylated proteins.

### Mass Spectrometry

Precipitated proteins were denatured and alkylated in 50 µl 8 M Urea, 1 M ammonium bicarbonate containing 10 mM tris (2-carboxyethyl) phosphine hydrochloride and 40 mM 2-chloro-acetamide. After 4-fold further dilution with 1 M ammonium bicarbonate and digestion with trypsin (250 ng/200 µl), peptides were separated from the beads and desalted with homemade C-18 stage tips (3 M, St Paul, MN). Peptides were eluted with 80% ACN and, after evaporation of the solvent in the speedvac, redissolved in buffer A (0,1% formic acid).

After separation on a 30 cm pico-tip column (75 µm ID, New Objective) in-house packed with C-18 material (1.9 µm aquapur gold, dr. Maisch) using a 140 minute gradient (5% to 80% ACN, 0.1% FA), peptides were delivered by an easy-nLC 1000 (Thermo) or easy-nLC1200 (Thermo), and directly electro-sprayed into an Orbitrap Fusion (Thermo Scientific), or an Orbitrap Exploris 480 (Thermo Scientific), or an Orbitrap Eclipse Tribrid Mass Spectrometer (Thermo Scientific). The Fusion was run in DDA mode with a cycle time of 1 second, MS1 scans were acquired in the Orbitrap over a m/z range of 400–1500 at a resolution of 240,000, with an AGC target of 75% and a 50 ms maximum injection time. Precursor ions for MS2 were selected using an intensity threshold of 15,000 ions, charge states of 2+ to 7+, and a dynamic exclusion of 60 s at 5 ppm. MS2 scans were performed in the ion trap at rapid resolution with a 1.6 m/z isolation window (using quadrupole isolation), using higher-energy collisional dissociation (HCD) at 30% collision energy, an AGC target of 100%, and a dynamic maximum injection time. The Orbitrap Exploris equipped with a FAIMS Pro interface, was run with a 1.5 s cycle time per FAIMS compensation voltage (CV), applying sequential CVs of –45 V and –65 V. MS1 scans were acquired over a m/z range of 375–1200 at a resolution of 120,000, with an AGC target of 300% and a 50 ms maximum injection time. For MS2, precursor ions were selected based on an intensity threshold of 5000 ions, charge states of 2+ to 6+, and a dynamic exclusion of 60 s at 10 ppm. Fragmentation was performed using higher-energy collisional dissociation (HCD) with 30% collision energy, a 0.7 m/z isolation window, a resolution of 7500, an AGC target of 150%, and a 30 ms maximum injection time. The Orbitrap Eclipse equipped with a FAIMS Pro interface was run with a 1.0 s cycle time per FAIMS compensation voltage (CV), applying sequential CVs of –45 V and –65 V. MS1 scans were acquired in the Orbitrap over a m/z range of 400–1400 at a resolution of 240,000, with an AGC target of 100% and a 50 ms maximum injection time. Precursor ions for MS2 were selected using an intensity threshold of 10,000 ions, charge states of 2+ to 7+, and a dynamic exclusion of 30 s at 10 ppm. MS2 scans were performed in the ion trap at rapid resolution with a 0.4 m/z isolation window (using quadrupole isolation), using higher-energy collisional dissociation (HCD) at 35% collision energy, an AGC target of 100%, and a dynamic maximum injection time.

The raw data was processed using MaxQuant (version 1.6.3.4) [116] with default settings: Oxidation (M) and Acetyl (Protein N-term) were set as variable modifications, and Carbamidomethyl (C) set as a static modification. Peptide length interval was set to 7-25 amino acids. The FASTA file used as the reference for the identification and quantification can be found in **Table S5** (available via https://doi.org/10.6084/m9.figshare.29967553). Label-free quantification (LFQ) was enabled, and iBAQ normalization was selected. The proteinGroups.txt output file of MaxQuant was used to follow up with data analysis, using a label-free quantification approach. For statistical analysis of MaxQuant output, either the Perseus framework (version 1.5.5.3.) offered by the Cox Lab [117], or R (version 4.0) was used for the analysis, following up the same workflow. Protein groups were filtered to remove contaminants and reverse hits, as well as, retaining only those with more than one unique peptide. iBAQ-normalized intensity values were log₂-transformed, and missing values were imputed using the minimum value approach.

To capture co-variation of streptavidin-enriched proteins between different BioID experiments (WT versus various 3xHA-TurboID strains) we performed principal component analysis (PCA) using total spectrum counts per protein present in each of the samples (MS/MS). Values for undetected proteins were set to 0. Values were ln(x + 1) transformed and PCA was performed and visualised using the ClustVis webserver (settings Nipals PCA, no scaling) [118]. Selection of principal components was based on those that accounted for the majority of the variance as well as showing clusters surrounding the specific TurboID-bait proteins (e.g. KiTT1,2 etc.).

### Cell cycle expression analysis

Cell cycle–resolved expression profiles were obtained from two published transcriptome datasets (RNA seq and microarray) of *T. thermophila* [51,52] (see **Table S3** for full list of proteins). Normalized expression values (Min-Max) were extracted for all candidate proteins identified by BioID (**Figure 2D**). KiTT1–4 and the majority of candidate kinetochore proteins showed a characteristic G2/M-enriched expression peak, consistent with their role in mitosis. Expression traces were plotted and clustered using custom R scripts v4.5. Clusters of expression traces were manually delineated based on average linkage clustering.

### Immunofluorescence

Cell were mixed with 0.5% Triton X-100 in PHEM buffer (60 mM Pipes, 25 mM HEPES, 2 mM MgCl_2_, 10 mM EGTA, pH6.9) for 20 sec and then fixed with 2% paraformaldehyde (Electron Microscopy science, Cat# 15710) in phosphate buffered saline (PBS) added on top of PHEM for 30 min at room temperature. The fixed cells were washed with PBS, mixed with Cultrex Reduced Growth Factor Basement Membrane Extract (BME) type 2 (R&D systems, Cat# 3533-010-02) in a 1:4 ratio of cells:BME, spread on slides and air-dried for 2 min. Slides were washed twice with PBS, permeabilized with 0.1% Triton X-100 in PBS (PBST) for 20 min, and blocked with 2% BSA in PBST for 1 h. The samples were incubated with primary antibodies diluted in 2% BSA PBST overnight at 4 °C. Slides were washed three times with PBS and incubated with secondary antibodies for 1 h at room temperature. After washing three times with PBS, slides were mounted with DAPI (Sigma-Aldrich, Cat# D9542) in ProLong Gold Antifade (Invitrogen, Cat# P36930).

### Image acquisition

Images were recorded with VT-iSIM imaging system (VisiTech) coupled to a Prime BSI Express sCMOS camera (Photometrix) on Nikon Ti2-E microscope. Imaging of Z-stacks with 0.2 µm intervals was performed with 100XC Sil 1.35-NA objective lens (Nikon). Images for **Figure 2B** and **Figure S5** were acquired with a Delta Vision Elite system (Applied Precision/GE Healthcare) using 100x/1.4-NA objective (Olympus) and deconvolved in SoftWorx (Applied Precision/GE Healthcare) software.

### RNAi image analysis

Maximum intensity Z-projection images were analyzed and quantified using the open-source software CellProfiler (version 4.2.6) [119]. Briefly, a pipeline was designed to first segment the MIC region based on the DAPI signal, and then within the MIC masksegment kinetochores based on HA signal. Average intensities of the HA and KiTT1^NDC80^ immunostainings were measured both in the segmented kinetochore regions as well as the non-kinetochore MIC region, which was taken as background. The average intensities of HA and KiTT1^NDC80^ in the background were subtracted to those obtained from the HA and KiTT1^NDC80^ signal in the kinetochore masks.

### Intra-kinetochore distance measurements

Images of metaphase micronuclei were processed using conservative deconvolution in Huygens Professional software (v20.10) and measured using Imaris software (10.0.1, Bitplane). First, the fluorescent objects were detected using the spot detection method with an estimated XY diameter of 150 nm and Z diameter of 300 nm. Sister kinetochore pairs were identified and the distance was measured between the corresponding proteins across these sister pairs (dKiTT1^NDC80^/dKiTT8^CENP-C^ and dHA/dmCherry). The distance between the centers of selected kinetochore pairs was obtained using the Measurement point tool. Only sister kinetochore pairs that can be detected as distinct, bi-oriented pairs were used for intra-kinetochore measurements. The dHA/dmCherry measurements were subtracted from the dKiTT1^NDC80^ measurement, or the inverse for dKiTT8^CENP-C^-based measurements. Then, the resultant number is divided by two to get the average intrakinetochore distance between HA/mCherry and KiTT1^NDC80^/KiTT8^CENP-C^, which is an effective strategy to account for chromatic aberrations in the imaging setup [67].

### Antibodies

Rabbit polyclonal antibodies were raised against residues 232-245 (C-SKHNSSKKNPQQNQ) of KiTT1^NDC80^ and residues 1017-1030 (C-MLKNTESQKYEKNE) of KiTT8^CENP-C^ (BioGenes GmbH). The specificity of the antibodies was verified using a peptide competition assay (**Figure S1**). The affinity-purified custom antibodies were used at 1:100 dilution. The CNA1 custom antibody had been developed previously and antiserum was used at 1:200 dilution (gift from Prof. Josef Loidl [35,120]). The commercial primary antibodies used were mouse anti-HA clone HA-7 (1:200, Sigma-Aldich, Cat# H3663), rabbit anti-HA (Sigma-Aldich, Cat# H6908), mouse anti-mCherry (4B3) (1:100, Invitrogen, Cat# MA5-32977), mouse anti-mNeonGreen (1:200 ChromoTek, Cat# 32f6), Streptavidin-488 (1:200, Invitrogen, Cat# S11223). Secondary antibodies were goat anti-rabbit conjugated to Alexa Fluor 488 (Invitrogen, Cat# A-11034), 568 (Invitrogen, Cat# A-11036), 647 (Invitrogen, Cat# A-21245), goat anti-mouse Alexa Fluor 488 (Invitrogen, Cat# A-11029), 568 (Invitrogen, Cat# A-11036), 647 (Invitrogen, Cat# A-21236) and used at 1:200.

## DATA AVAILABILITY

All data needed to evaluate the conclusions in the paper are present in the paper and/or the supplementary materials, and via Figshare: https://doi.org/10.6084/m9.figshare.29967553. Mass spectrometry proteomics data have been deposited to the ProteomeXchange Consortium via the PRIDE [121] partner repository with the dataset identifier PXD070467. Relevant constructs and strains will become available through the Tetrahymena Stock Centre.

## ETHICS DECLARATIONS

The authors declare no conflicts of interests.

## AUTHOR CONTRIBUTIONS

**EIA:** Writing–original draft; visualization; formal analysis; investigation; data curation; writing-review and editing; methodology; conceptualization; resources**. MWDR:** Writing–original draft; visualization; formal analysis; investigation; data curation; writing-review and editing; methodology; conceptualization; resources. **LEvR:** Formal analysis; investigation; data curation; writing-review and editing; resources; conceptualization. **PSA:** Resources; data curation; formal analysis. **HRV:** Resources; data curation; formal analysis**. ECT:** Writing–original draft;visualization;formal analysis;investigation;data curation; writing-review and editing; methodology; conceptualization; resources; supervision; project administration; funding acquisition. **BS:** Writing–original draft;visualization;formal analysis;investigation;data curation; writing-review and editing; methodology; conceptualization; resources; supervision; project administration; funding acquisition. **GJKPL:** Writing–original draft;visualization;formal analysis;investigation;data curation; writing-review and editing; methodology; conceptualization; resources; supervision; project administration; funding acquisition

## ACKNOWLEDGEMENTS AND FUNDING

This work was funded by the Hubrecht Institute/KNAW and the European Research Council (ERC-SyG 855158). This work is part of the research programme VICI with project number 016.160.638, which is (partly) financed by the Netherlands Organisation for Scientific Research (NWO). E.T. was supported by an NWO-VENI Fellowship (VI.Veni.202.223). We thank Prof. Josef Loidl, Dr. Rachel Howard-Till (University of California Davis, Davis, CA, USA), Dr. Miao Tian (Ocean University of China, Qingdao, China), Dr. Kazufumi Mochizuki (IGH, CNRS - Université de Montpellier, Montpellier, France) for the cell lines, tagging constructs and CNA1 antibody. We thank Dr. Lev Tsypin (Stanford University, USA) for sharing re-normalized values for cell cycle expression data ahead of publication. We thank Dr. Jie Xiong (Institute of hydrobiology, Wuhan, Haibei, China) for sharing the full predicted proteomes of 10 Tetrahymena species underlying the online blast server at http://ciliates.ihb.ac.cn/. We thank Tessa Alofs for maintaining cell culture, Dr. Victor Tiroille for assistance with CellProfiler software, Dr. Carlos Sacristan for assistance with data analysis, and Noa Mutsters and Tessa Verschuuren for assisting with tagging and micronuclei sample preparation. We thank the Hubrecht Imaging Center for assistance with all imaging experiments. We thank Evelien Stouten and Jolanda Schuurmans (Utrecht University, Utrecht, Netherlands) for their assistance with the biolistic gun, and Joris Treep for preliminary work on this project. We thank all members of the Snel and Kops groups for discussion and comments on the manuscript.

## SUPPLEMENTARY DATA

Supplementary data for figures 2A, 3, 4B, 5C, 6 and 7, is accessible via https://doi.org/10.6084/m9.figshare.29967553.

## SUPPLEMENTARY TABLES

***Table S1. HHsearch output for searches using profile HMMs of KiTTs against a database of profile HMMs of kinetochore components and COG/KOG database***

Includes a summary Table of −log10 (E) values for HMM-vs-HMM searches (HHsearch) and separate sheets for searches per KiTTs (in blast tab output format).

***Table S2. LC-MS/MS data for BioID experiments***

Overview of the mass spectrometry data for all BioID experiments used in this study.

***Table S3. Cell Cycle Expression***

Data underlying Figure 2E with cell cycle expression analysis using RNA seq and microarray data adapted from Bertagna *et al.* [51].

***Table S4. oligos and plasmids***

Overview of oligos and plasmids used in this study.

***Table S5. Tetrahymena thermophila predicted proteome version used in this study***

Includes two tabs: (1) full table of the predicted proteome and associated annotation as found in Uniprot - downloaded on 3 march 2024 - https://www.uniprot.org/proteomes/UP000009168. (2) conversion table for TTHERM IDs associated with the Uniprot proteome (2016 and 2020 predicted proteome versions) and the new 2024 predicted proteome.

## SUPPLEMENTARY FIGURES

**Figure S1.**
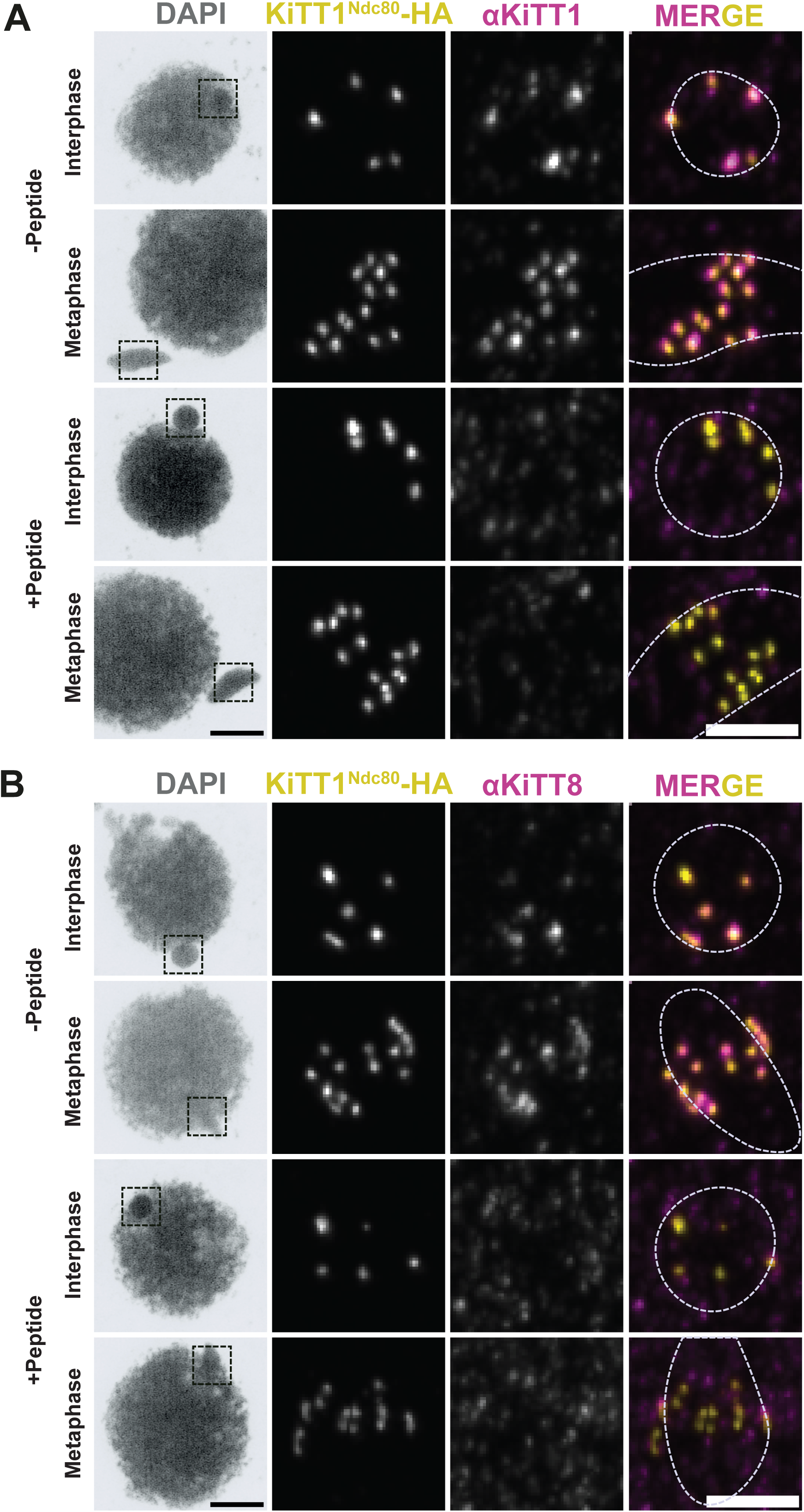
Validation of the specificity of KiTT1^Ndc80^ and KiTT8^CenpC^ antibodies using a peptide competition assay. KiTT1^NDC80^ and KiTT8^CENP-C^ antibodies were pre-incubated with a five-fold excess of synthetic peptides corresponding to 232-245 residues of KiTT1^NDC80^ and 1017-1030 residues of KiTT8^CENP-C^ respectively. Cells expressing HA3-TurboID tagged KiTT1^NDC80^ were fixed and stained with DAPI, anti-HA (yellow), (A) anti-KiTT1^NDC80^, peptide blocked KiTT1^NDC80^ antibody (magenta), (B) anti-KiTT8^CENP-C^, peptide blocked KiTT8^CENP-C^ antibody (magenta). MIC outline is shown with the dotted white line. Scale bars 5 µm, insets 2 µm.

**Figure S2.**
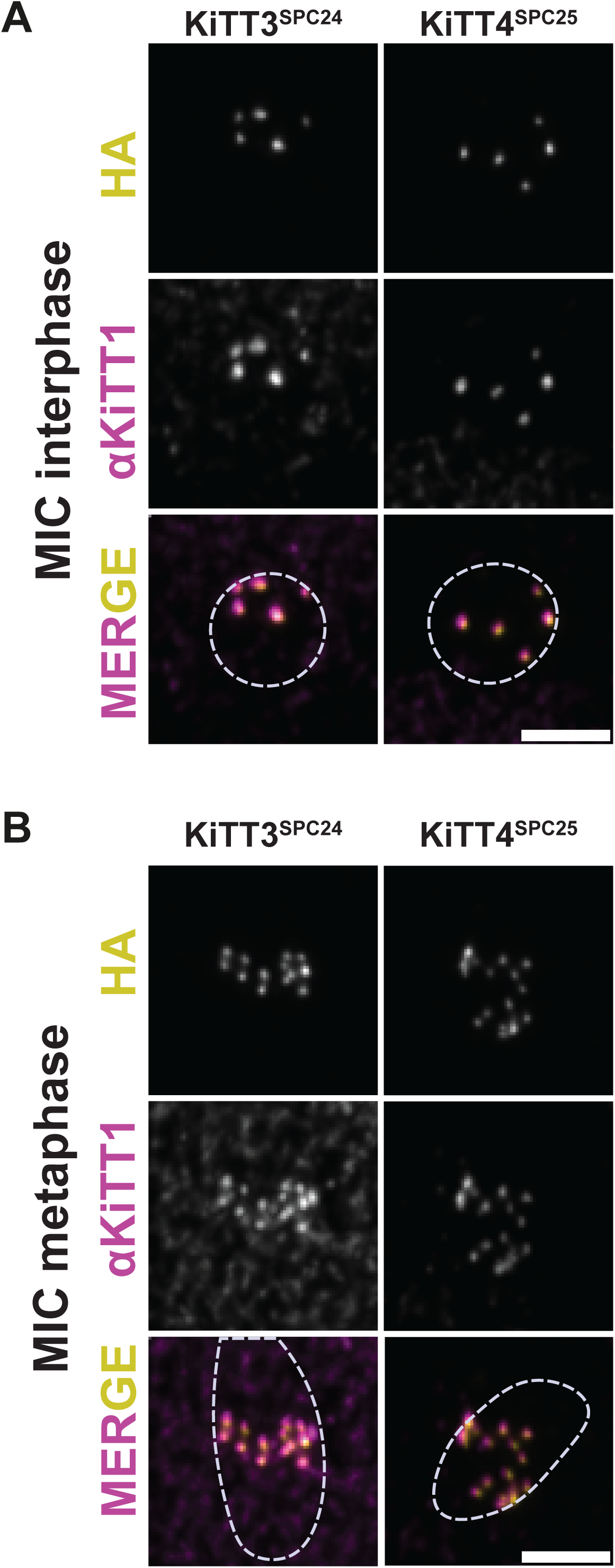
Co-staining of putative orthologs of KiTT3(SPC24) and KiTT4(SPC25) and αNDC80. Immunofluorescence staining of interphase (**A**) and metaphase (**B**) MICs stained with antibodies against KiTT1^NDC80^ (magenta) and HA-tagged putative orthologs of SPC24 (TTHERM_00312495) and SPC25 (TTHERM_01142690; yellow). Scale bar, 2 µm.

**Figure S3.**
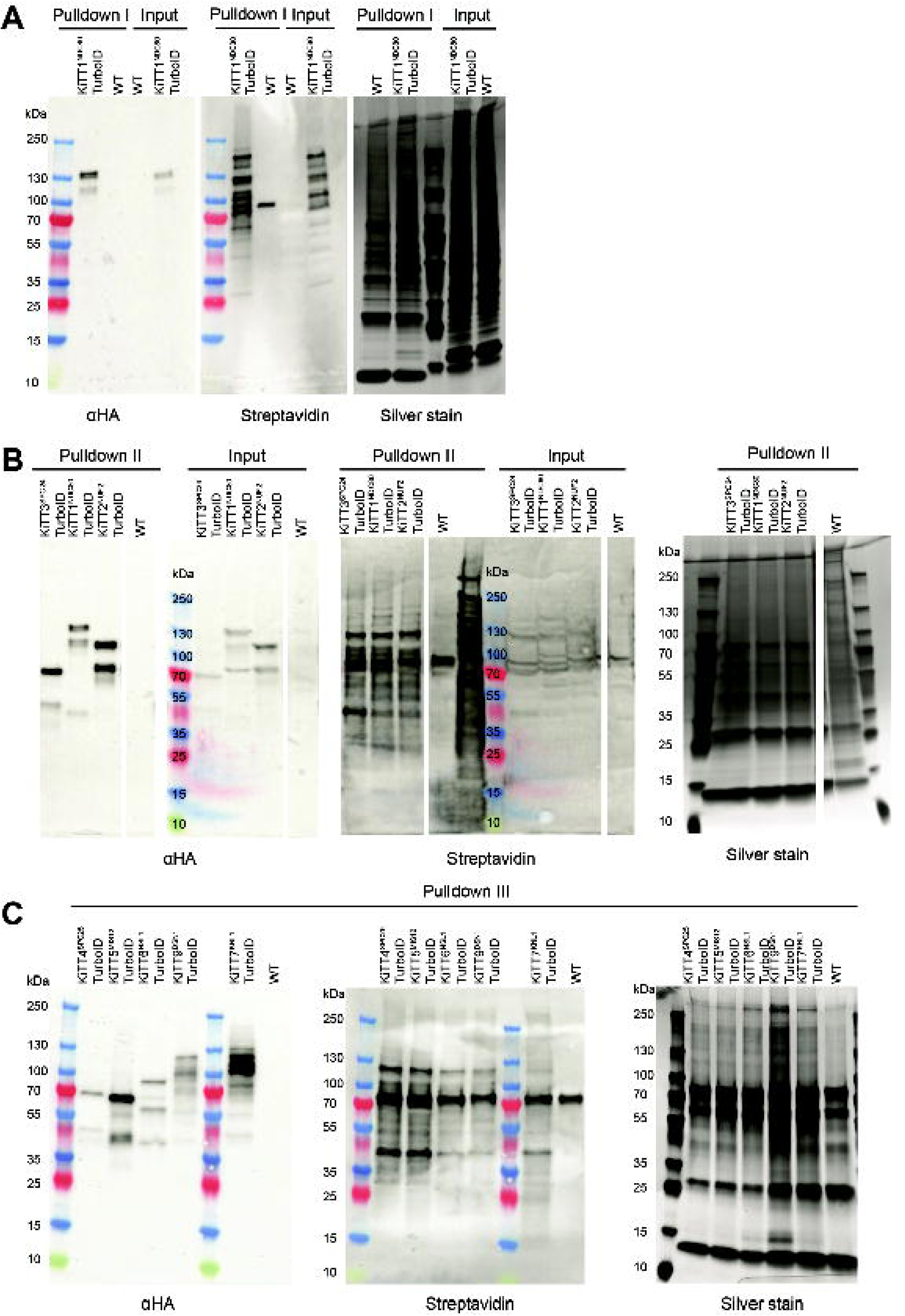
Pulldown of biotinylated proteins. Western blot of affinity purified proteins from control and cells expressing HA3TurboID tagged bait proteins. (A) Pulldown of 3xHA-TurboID-tagged KiTT1^Ndc80^ cells. The left-hand blot shows the staining for HA, showing bands corresponding to the 3xHA-TurboID-tagged KiTT1^Ndc80^. Middle, staining for streptavidin, showing that the 3xHA-TurboID-tag results in robust biotinylation of a range of proteins. Right, a control for total protein in the lysate using a silver stain. (B) Same as **A** but here showing the results obtained from cells with 3xHA-TurboID-tags on KiTT1^Ndc80^, KiTT2^NUF2^, KiTT3^SPC24^ (**C**) Same as **A** showing the results obtained from cells with 3xHA-TurboID-tags on KiTT4^SPC25^, KiTT5^MIS12^, KiTT6^NSL1^, KiTT7^KNL1^, KiTT9^DSN1^.

**Figure S4.**
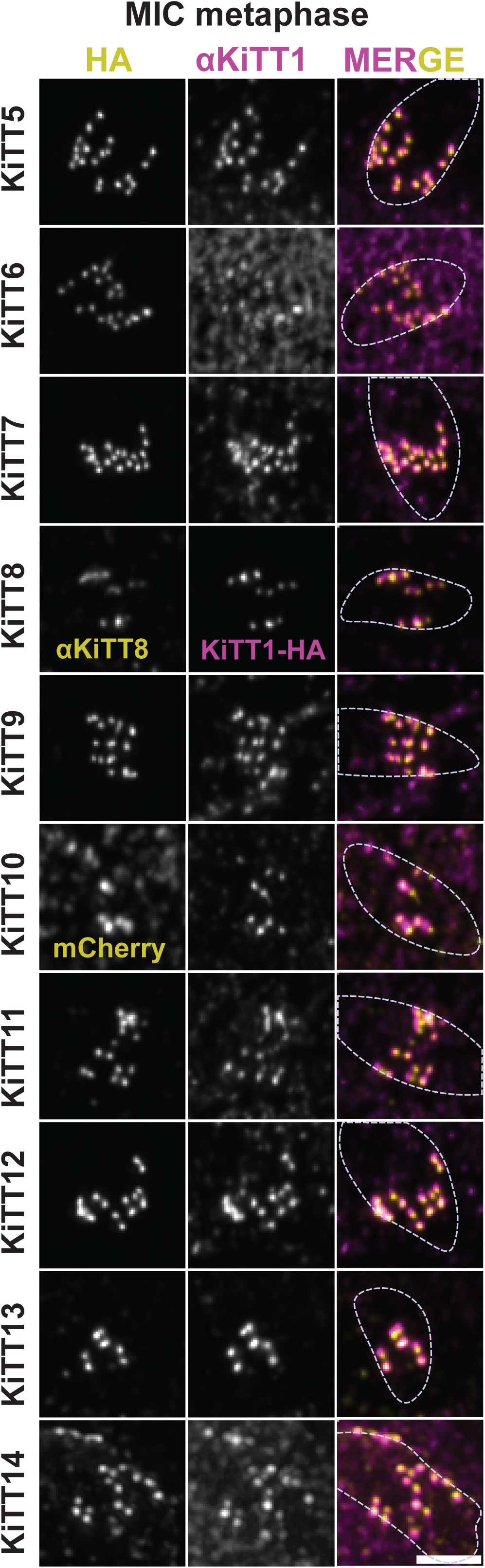
Immunofluorescence staining of metaphase MICs. The cells expressing 3xHA-TurboID-tagged candidates were stained with antibodies against HA and KiTT1^NDC80^. KiTT8 was detected with a custom antibody against KiTT8 in KiTT1^NDC80^-HA3TurboID cells. KiTT10-mCherry cells were co-stained with mCherry and KiTT1^NDC80^ antibodies. Scale bar, 2 µm.

**Figure S5.**
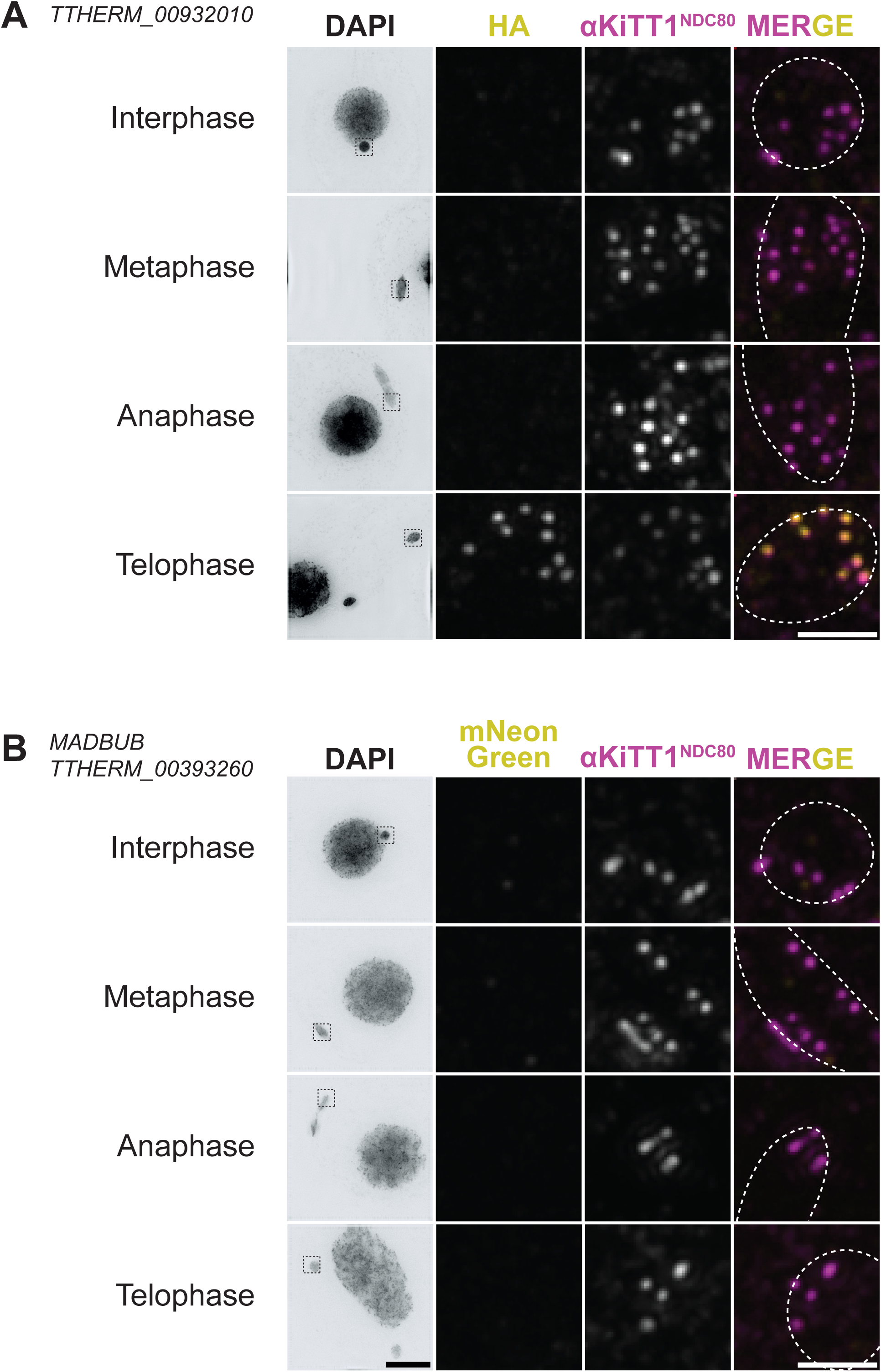
Localisation of TTHERM_00932010 and TTHERM_00393260^MADBUB^ across interphase and different stages of MIC mitosis. Immunostaining of TTHERM_00932010-3xHA-TurboID (**A**) or TTHERM_00393260^MADBUB^ (**B**) and KiTT1^NDC80^ in interphase and during mitosis. In the merged images, TTHERM_00932010-3xHA-TurboID/TTHERM_00393260^MADBUB^-3xHA-TurboID is shown in yellow and KiTT1^NDC80^ in magenta. Images were acquired using the Delta Vision Elite system. Scale bar 10 µm, insets 2 µm.

**Figure S6.**
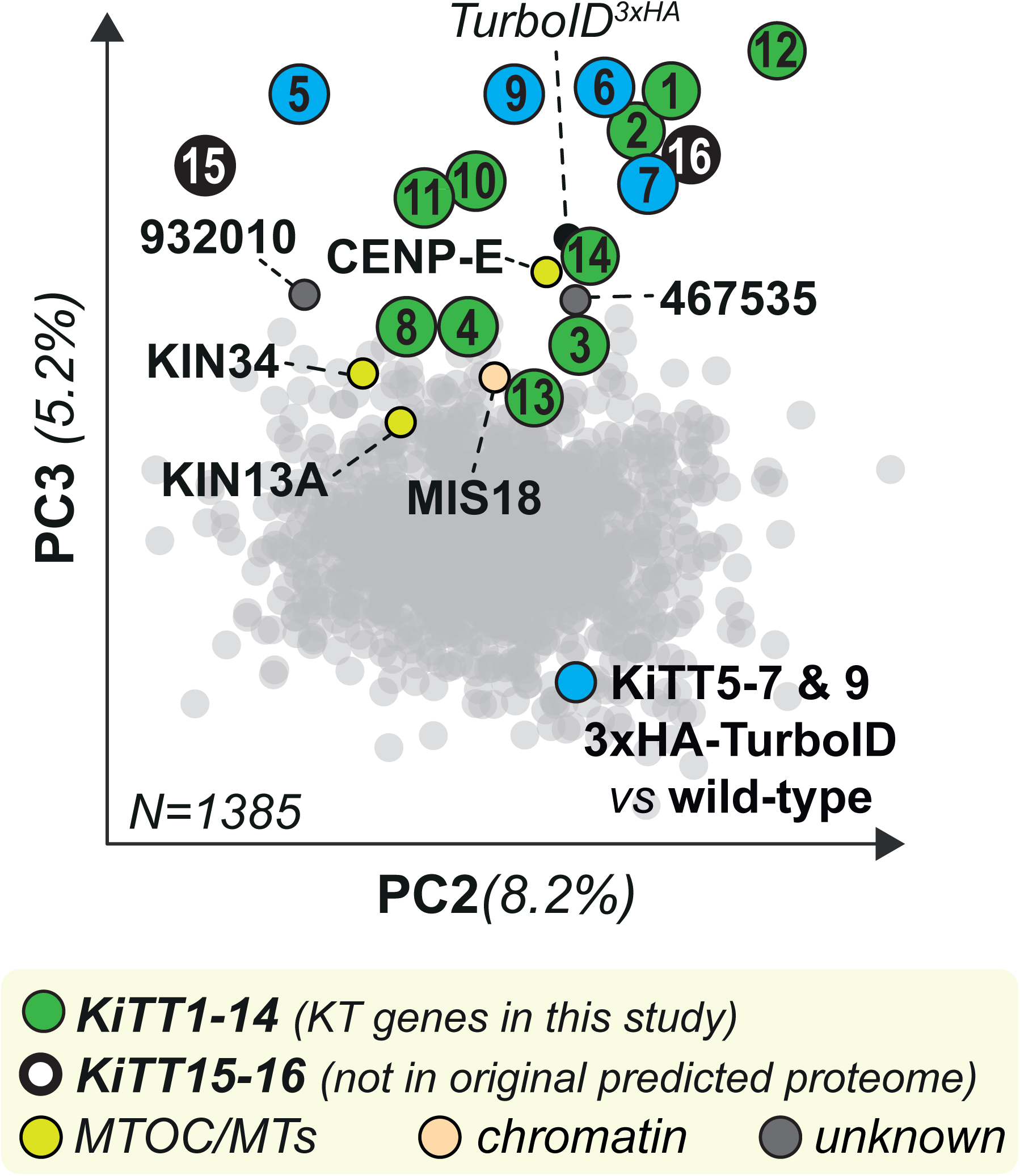
BioID for KiTT5-7 & KiTT9. PCA of integrated spectral counts detected in BioID experiments from KiTT5^MIS12^, KiTT6^NSL1^, KiTT7^KNL1^, KiTT9^DSN1^ and wildtype cells as a control (see **Table S2** for MS analysis). PC2 and PC3 are plotted as they showed a clear kinetochore-based cluster, similar to Figure 2D. MIS12 complex proteins (+ KNL1) are highlighted in blue, characterized KiTTs (5-14) in green and in black KiTT15-16 that were not part of the original proteome that was used to analyse the mass spectra. Other categories have unique colors (MTOC/SPB & chromatin) and unknown proteins are grey.

**Figure S7.**
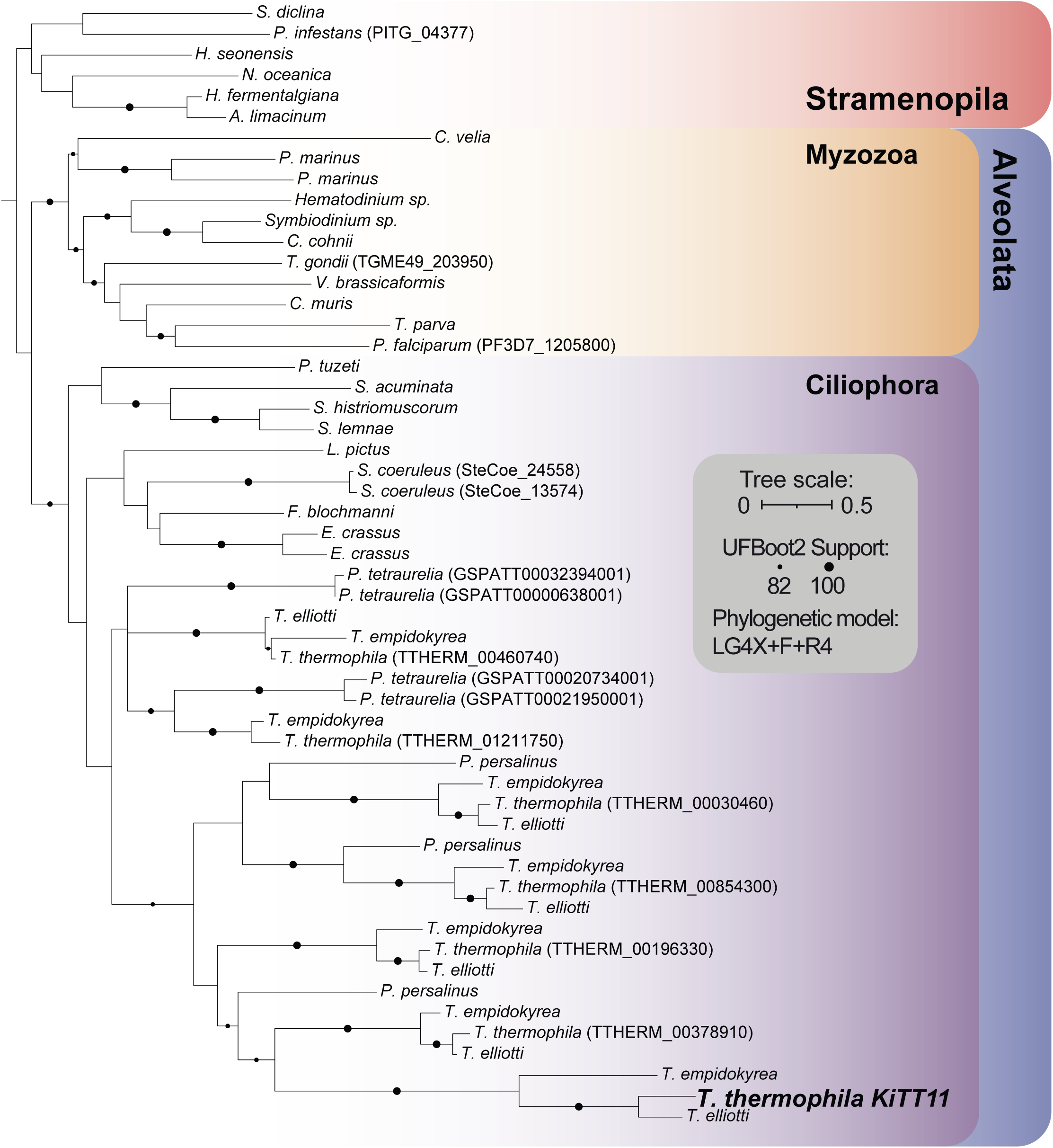
Phylogenetic tree of Bdp1 homologs across stramenopiles and alveolates. Phylogenetic tree of the SANT domain of KiTT11 and Bdp1 orthologs across Stramenopila and Alveolata. KiTT11 arose from a *Tetrahymena-*specific duplication. Bdp1 orthologs were obtained using homology searches in a database derived from [126]. A maximum-likelihood was run using IQ-TREE with the LG4X+F+R4 substitution model. Black dots show nodes with ≥ 82 UFBootstrap2 support, as specified in in-figure legend.

**Figure S8.**
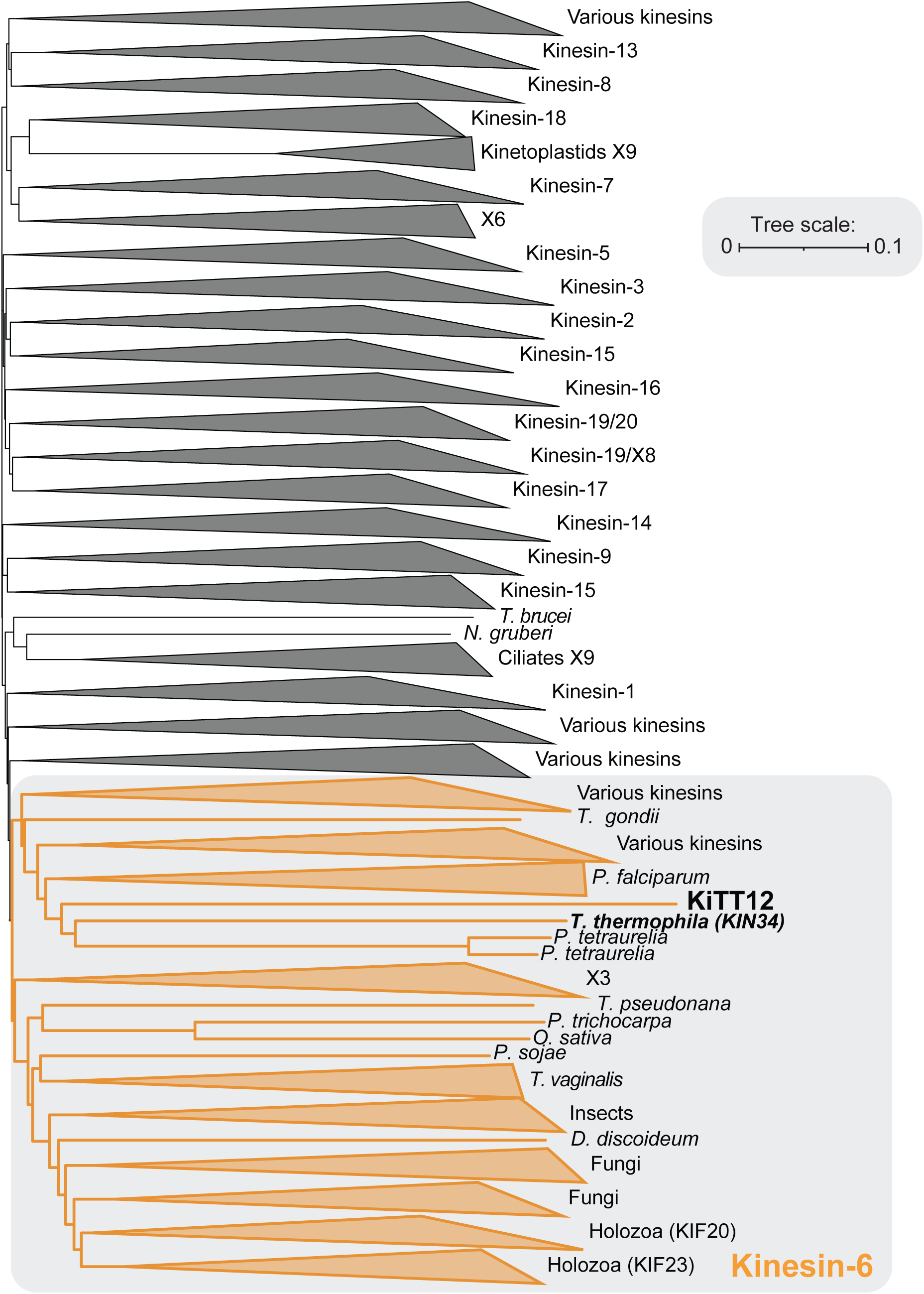
Structure-based phylogeny of kinesin motor domain folds across eukaryotes. Phylogenetic tree inferred using Foldtree [111] of the predicted structures of kinesin motor domains, predicted from the sequences from Wickstead et al [74], with additionally the predicted motor domain fold of KiTT12 (highlighted in bold). An interactive version of this tree can be accessed via https://itol.embl.de/tree/18522192164228841760699518.

**Figure S9.**
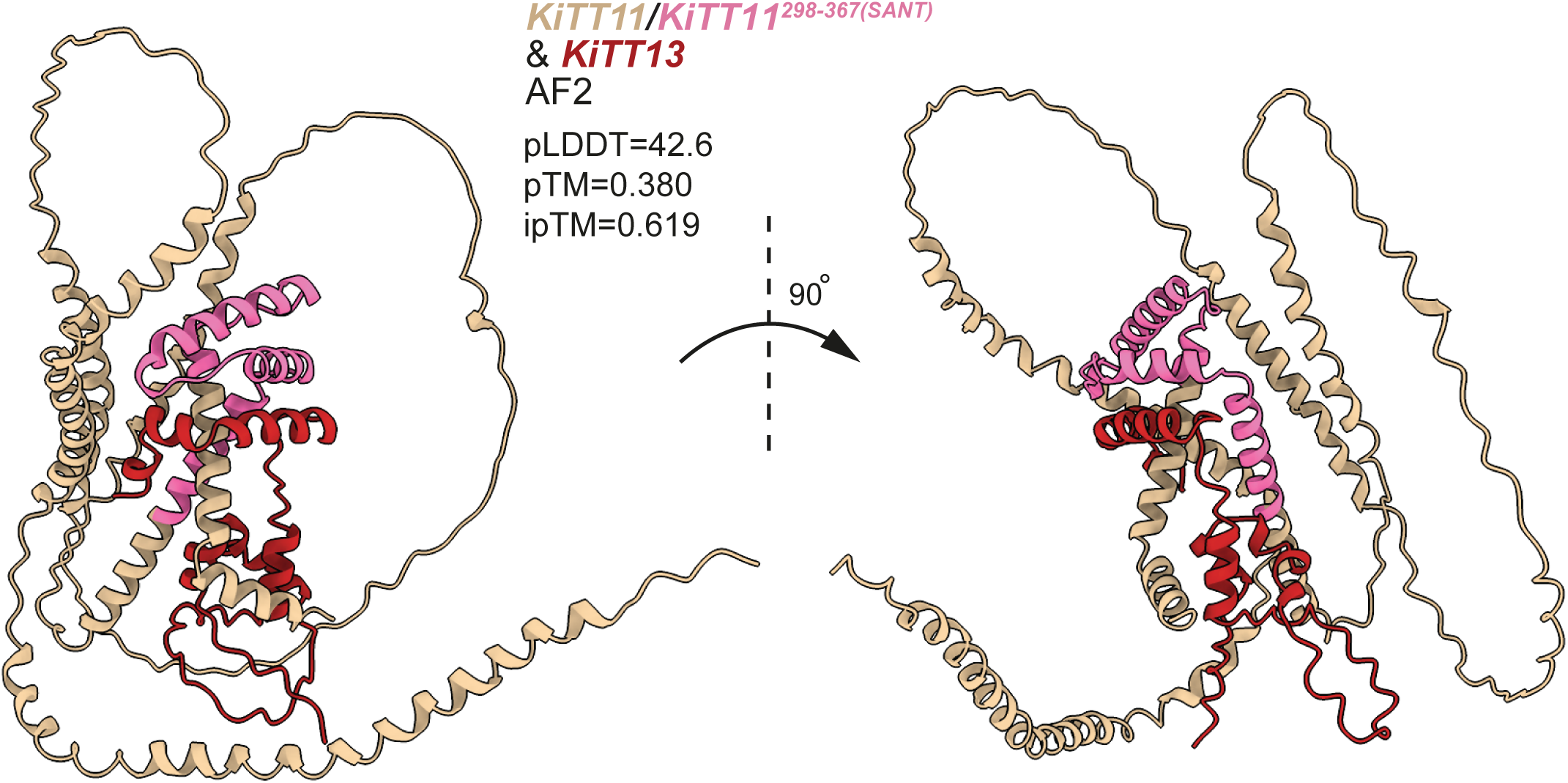
AF2 co-fold prediction of KiTT11 and KiTT13. KiTT11 is shown in tan with the SANT domain in pink. KiTT13 is shown in red. Right shows a 90° rotation perpendicular to the plane of the figure.

